# The molecular structural basis of the braking action of muscle

**DOI:** 10.64898/2026.09.25.753934

**Authors:** Cameron Hill, Michaeljohn Kalakoutis, Alice Arcidiacono, Atsuki Fukutani, Theyencheri Narayanan, Elisabetta Brunello, Luca Fusi, Malcolm Irving

## Abstract

Skeletal muscles generate force and shortening through a structural ‘powerstroke’ in actin-bound myosin motors linked to ATP hydrolysis, but they also act as brakes to resist stretching by an external load. The molecular structural basis of muscle braking has remained obscure. Here we show that muscle braking is driven both by structural changes in myosin motors and by distortion of the lattice of thick and thin filaments including their M-band and Z-disk connections. A strained motor conformation is established early during stretch, but force continues to increase as thin filaments are displaced from their symmetrical lattice positions, the lateral shear between thick filaments increases, and the lattice volume decreases. Moreover, whilst the motor conformation rapidly recovers after the stretch, the distortion of the filament lattice is largely maintained, storing the work done on the muscle during stretch and accounting for the residual force enhancement after stretch.

## Introduction

Skeletal muscles are often considered as biological machines for converting chemical energy into mechanical work, such as when lifting a heavy object. However, muscles also act as brakes to resist an external force whilst being stretched, whilst walking down stairs, for example. In this stretching or braking mode, mechanical work is done *on* the muscle as it is extended. In animal locomotion, muscles typically alternate between shortening and braking modes, and the work done on muscles and tendons during each braking phase is stored for release in the next shortening phase, increasing the overall efficiency of locomotion^1^.

Research into the molecular basis of muscle function has been largely focused on its shortening mode, which is driven by the relative sliding between the myosin-containing thick filaments and actin-containing thin filaments. Filament sliding during shortening, in turn, is driven by a cycle in which the motor domain of myosin attaches to an actin site in the thin filament, changes conformation in a molecular ‘powerstroke’ and then detaches, in a biochemical cycle linked to ATP hydrolysis^2–6^. Detailed structural and mechanical models based on these concepts have been tested in intact muscles and isolated muscle fibres by *in situ* structural methods like time-resolved X-ray diffraction. The results of such studies on shortening muscle provided detailed support for molecular models based on the powerstroke paradigm^7–10^.

The molecular mechanisms underlying the braking mode of muscle function are less well understood. Muscle sarcomeres remain stiff during braking^11^ showing that myosin motors are bound to thin filaments, but their cycle of interaction with actin and ATP is fundamentally different from that observed during shortening. The mechanical work done on the muscle during braking is larger than the heat produced^12,13^, showing that muscles absorb energy during braking. The stored energy is too large to be accounted for by the strain of actin-attached myosin motors^13^.

Mechanical studies of muscle braking in different muscle types and preparation showed that muscle force remains higher for several seconds after a stretch, a phenomenon referred to as residual force enhancement^14,15^. Moreover, part of the force increase during a stretch and most of the force enhancement after stretch seems to be associated with structures other than the myosin motors^16,17^, including the sarcomeric cytoskeleton^18^, which organises the lattice of thin and thick filaments through the cross-linking components of the M-band, Z-disk and titin. Different cytoskeletal structures may be involved at different sarcomere lengths or at different speeds of stretch. Inhomogeneity of sarcomere lengths along the muscle fibres may also be involved in the braking response in more extreme conditions.

Here we used small-angle X-ray diffraction (SAXD) and X-ray interference to investigate both the myosin-related and the cytoskeletal components of the braking action of intact extensor digitorum longus (EDL) muscles of the mouse. We used ultra small-angle X-ray diffraction to characterise changes in the sarcomere length distribution along the same muscles during braking. The results allowed us to determine the relative contributions of myosin and cytoskeletal structures to force enhancement during muscle braking in conditions in which sarcomere length dispersion is negligible. They reveal the time course of each contribution, both during and after the applied stretch. Finally, they provide a structural description of the conformational changes in actin-attached myosin head domains during braking, and of the distortions of the lattice of thick and thin filaments that could store energy during stretch.

## Results

### Force and sarcomere length

The length of EDL muscles isolated from the mouse was adjusted to give a mean sarcomere length of 2.56 ± 0.01 µm (mean ± SE, n=8) as determined from the position of the first-order X-ray reflection generated by the sarcomeres (Table S1; Fig. 1a,b,c, cyan). Muscles were then stimulated repetitively at 27°C (Fig. 1a). Muscle length was initially held constant as fixed-end force developed to 245 kPa and sarcomeres shortened to 2.31 µm (Fig. 1b,c, red) as the tendons were extended. Sarcomere length dispersion (σSL) determined from the axial width of the first-order sarcomere reflection was about 0.11 µm, both at rest and at peak force before the stretch (Fig. 1d).

**Fig. 1.**
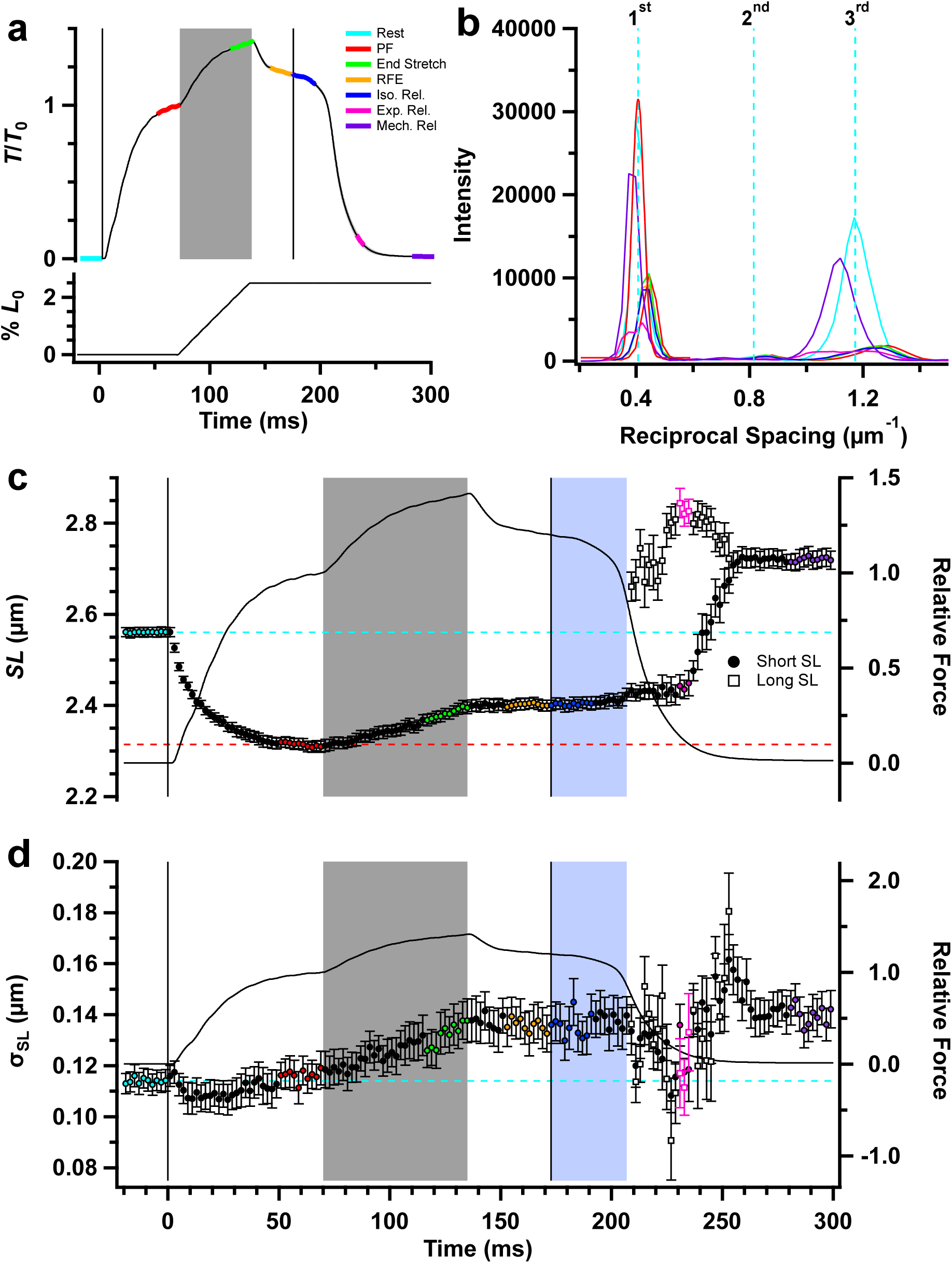
Force and sarcomere length changes in tetani with ramp stretch. **a** Force (*T*/*T*_0_) normalised with respect to the average force in the 10 ms time period before a constant velocity stretch of 2.5% *L*_0_ (grey shading). Coloured segments of the force trace denote periods used for averaging the X-ray data: cyan, rest; red, peak force (PF) before the stretch; green, end of ramp stretch; orange, residual force enhancement (RFE) after stretch; blue, isometric relaxation; magenta, late exponential relaxation when *T*/*T*_0_ is ∼10%; purple, mechanically relaxed. Mean ± S.E.M. for n = 7 muscles. **b** Ultra-small-angle meridional intensity distribution showing sarcomere reflections collected during the identified time periods after subtraction of the diffuse background, with orders of sarcomere reflections labelled. **c** Sarcomere length (*SL*). **d** Sarcomere length dispersion (σ*SL*). Data in **c** and **d** were determined from single Gaussian fits of the first-order sarcomere reflection during isometric relaxation and double-Gaussian fits during exponential relaxation, when the shorter and longer sarcomere populations are denoted by filled circles and open squares respectively. The average force from **a** is superimposed on the sarcomere length data in **c** and **d**, and horizontal dashed lines denote the average resting (cyan) and pre-stretch peak force value (red) of each parameter. Colour coding in **b**-**d** denotes periods shown in **a**. Vertical solid lines in **a**, **c** and **d** denote time of delivery of the first (*t* = 0 ms) and final (*t* = 173 ms) electrical stimulus. The blue shaded region denotes isometric relaxation. Data in **c**-**d** are mean ± S.E.M. for 15 records from 7 muscles.

After force had developed in fixed-end conditions, the muscles were stretched by 2.5% muscle length at constant velocity (Fig. 1, grey shading). Mean sarcomere length in the centre of the muscle increased by 0.08 µm (Fig. 1c, green), corresponding to filament sliding of about 40 nm in each half-sarcomere, which is expected to be large enough to detach all the myosin motors that were attached to actin before the stretch. Force increased by about 40% during the stretch; after the stretch it declined rapidly for about 10 ms, then much more slowly. At the time of the last stimulus force was still 22% higher than that before the stretch. Sarcomere length did not change significantly during this period of residual force enhancement (Fig. 1b,c; green, orange). σ_SL_ increased slightly to 0.13 µm during the stretch and during residual force enhancement (Fig. 1d, green, orange).

After the last stimulus, force, sarcomere length and sarcomere dispersion were well maintained during isometric relaxation (Fig. 1a,c,d; blue shading). Then, about 35 ms after the last stimulus and coinciding with an inflection point in the force trace, two populations of sarcomeres with different lengths became visible (Fig. 1c). The later force relaxation was well-described by a single exponential function with rate constant 81.0 ± 4.7 s^−1^. The shorter population of sarcomeres (filled circles) retained a length of about 2.4 µm for the first 30 ms of exponential relaxation, while the longer population were stretched to about 2.8 µm. Subsequently, the lengths of the two populations converged, and by 75 ms after the last stimulus only a single population could be detected with sarcomere length 2.72 ± 0.02 µm (Fig 1c).

### The lattice of thick and thin filaments

The hexagonal lattice of thick and thin filaments produces a series of equatorial X-ray reflections (Fig. S1a), of which the strongest are the 1,0 reflection, associated with the planes passing through the centres of the thick filaments bordering the unit cell, and the 1,1 reflection, associated with the planes through the centres of both thick and thin filaments (Fig. 2c, inset). The intensity of the 1,0 reflection (*I*_1,0_) decreased during fixed-end contraction (Fig. 2a), and that of the 1,1 reflection (*I*_1,1_) increased (Fig. 2b). These reciprocal changes are mainly due to movement of myosin motors away from the folded conformation on the surface of the thick filaments that they occupy in resting muscle, to an actin-attached conformation during fixed-end contraction in which more of their mass is associated with the thin filaments^19^, although these X-ray reflections are also sensitive to sarcomere length and disorder of the filament lattice.

**Fig. 2.**
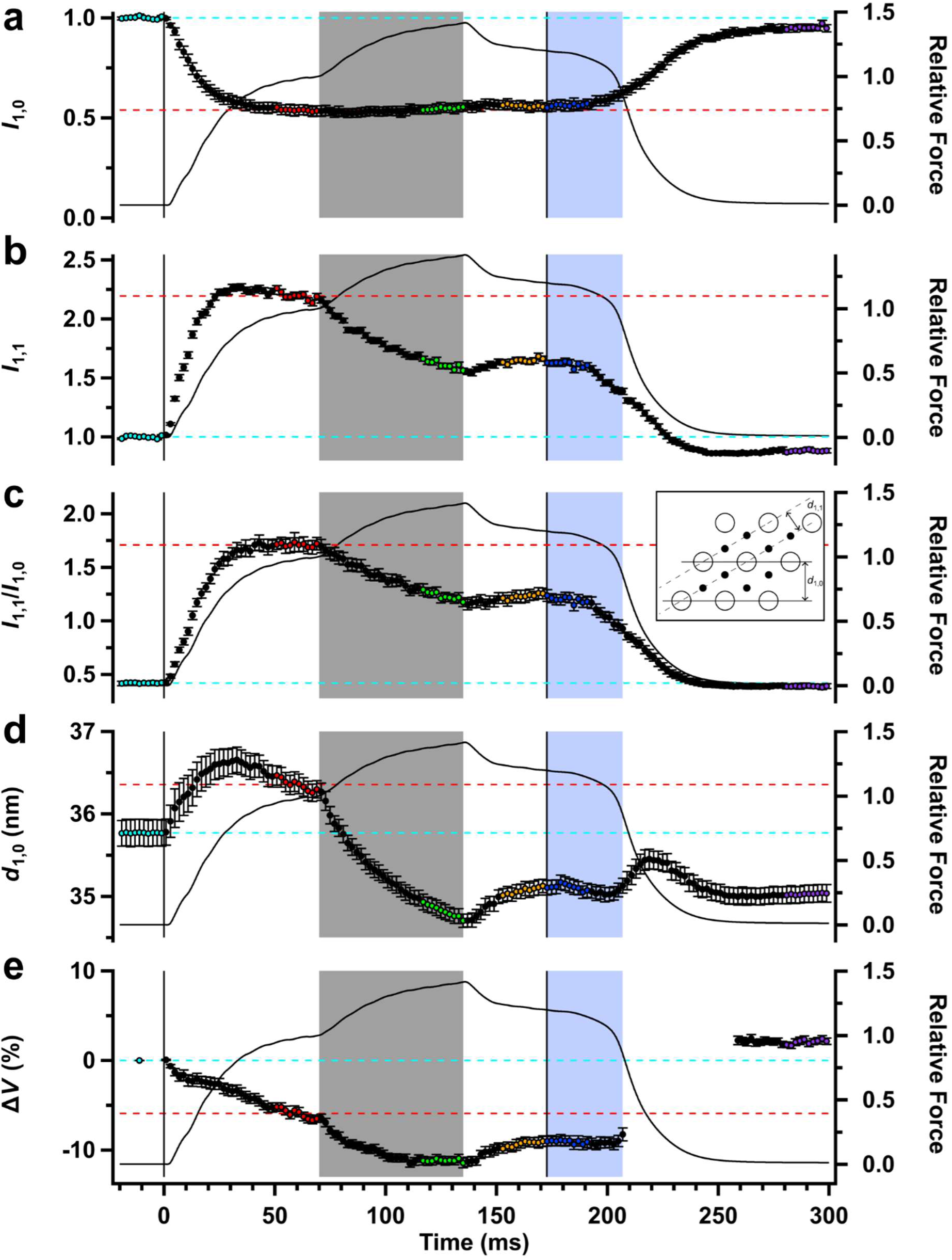
Equatorial X-ray reflections. **a** Intensity of the 1,0 reflection (*I*_1,0_). **b** Intensity of the 1,1 reflection (*I*_1,1_). **c** Equatorial intensity ratio (*I*_1,1_/*I*_1,0_). *Inset*: hexagonal filament lattice showing the 1,0 and 1,1 planes and their spacings *d*_1,0_ and *d*_1,1_, respectively. **d** Spacing of the 1,0 lattice planes (*d*_1,0_). **e** Volume of the filament lattice (Δ*V*) expressed as a percentage change from the resting value. Relative force (*T*/*T*_0_) is shown as the continuous black line. Colour coding of data points as in Fig. 1. Vertical lines denote the start and end of electrical stimulation; horizontal dashed lines denote the average resting value (cyan) and pre-stretch peak force value (red) of each parameter. Mean ± S.E.M. for n = 8 muscles.

The changes in *I*1,0 and *I*1,1 during the stretch were strikingly non-reciprocal, as first reported in frog muscle^20^. *I*_1,0_ did not change during the stretch (Fig. 2a), but *I*1,1 decreased by about 30% (Fig. 2b), suggesting that the stretch displaced thin filaments from their trigonal positions in the filament lattice (see Discussion). *I*_1,1_ recovered slightly as stimulation continued after the stretch but remained below its pre-stretch value. *I*_1,0_ and *I*_1,1_ were almost constant for about 20 ms after the last stimulus, then started to recover reciprocally towards their resting values.

The radial widths of the 1,0 and 1,1 equatorial reflections, σ_1,0_ and σ_1,1_ respectively, increased substantially during fixed-end force development (Fig. S2a), signalling an increase in the disorder of the filament lattice^21^. There was no change in σ1,0 during the stretch, but σ_1,1_ increased by about 4%. For most of the protocol, the ratio σ_1,1_/σ_1,0_ (Fig. S2b) was close to the theoretical value of √3 expected for dispersion of filament lattice spacings in the absence of paracrystalline or liquid-like lattice disorder, although there were deviations from this value at the start of stimulation and during exponential relaxation.

The spacing of the filament lattice, measured as the separation of its 1,0 planes (*d*1,0, Fig. 2d), increased slightly during fixed-end force development, and the volume of the filament lattice (Fig. 2e) decreased by about 6% because the lattice expansion does not completely compensate for the sarcomere shortening, as reported previously^22,23^. *d*_1,0_ decreased further during the stretch and partially recovered after the stretch, so that its time course mirrored that of force rather than sarcomere length. Filament lattice volume decreased during the stretch, becoming about 11% smaller than at rest, and recovered only partly during the residual force enhancement. These changes indicate that the filament lattice is compressed by the stretch, potentially storing energy in the sarcomeric cytoskeleton, and that the compression is not fully reversed during residual force enhancement.

### Thick filament regulation

In resting muscle, myosin motors are folded back against their tails on the surface of the thick filaments in a roughly helical arrangement with a 43 nm periodicity, making them unavailable for actin interaction and ATP hydrolysis. During contraction, motors are released from this folded inhibited state and become available for actin binding. The folded helical state of the myosin motors can be monitored in intact muscles using the myosin-based layer line X-ray reflections. The first-order myosin-based layer line (ML1) is the brightest of these layer lines (Fig. S1c). Its intensity (*I*_ML1_) decreased to about 10% of the resting value during fixed-end force development (Fig. 3a; Table 1) but there was no further change in *I*_ML1_ either during or after the stretch. The same pattern was observed in the intensity of the fourth myosin-based layer line (*I*_ML4_; Fig. 3b). We conclude that the regulatory state of the thick filament is not altered by the stretch or the accompanying increase in force.

**Fig. 3.**
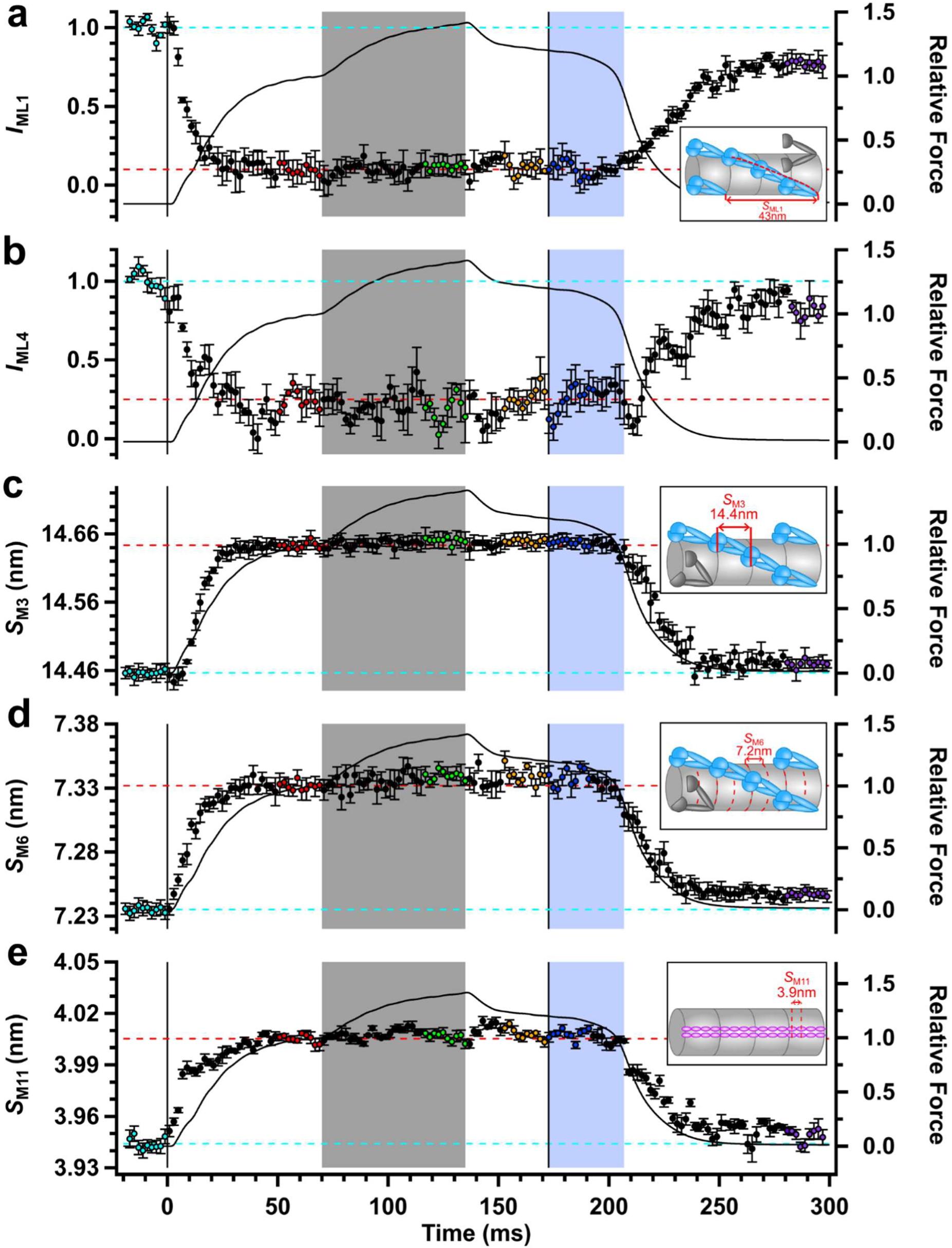
X-ray signals associated with thick filament activation. **a** Intensity of the first myosin layer line (*I*_ML1_). *Inset*: Helical arrangement of pairs of myosin motors (blue) folded back against their tails that gives rise to the ML1 reflection with axial periodicity 43 nm; disordered motors are shown in grey. **b** Intensity of the fourth myosin layer line (*I*_ML4_). **c** Spacing of the third myosin-based meridional reflection (*S*_M3_) corresponding to the axial periodicity of about 14.4 nm in resting muscle shown in the *Inset*. **d** Spacing of the sixth myosin-based meridional reflection (*S*_M6_) corresponding to the axial periodicity of about 7.2 nm in resting muscle shown in the *Inset*. **e** Spacing of the eleventh myosin-based meridional reflection (*S*_M11_) corresponding to the 3.9 nm axial periodicity of titin domains in the thick filament C-zone shown in the *Inset* as magenta ellipses. Relative force (*T*/*T*_0_) is shown as the continuous black line. Colour coding of data points as in Fig. 1. Vertical lines denote the start and end of electrical stimulation; horizontal dashed lines denote the average resting value (cyan) and pre-stretch peak force value (red) of each parameter. Mean ± S.E.M. for n = 5 muscles in **a** to **d**, and n = 3 muscles in **e**.

**Table 1.**
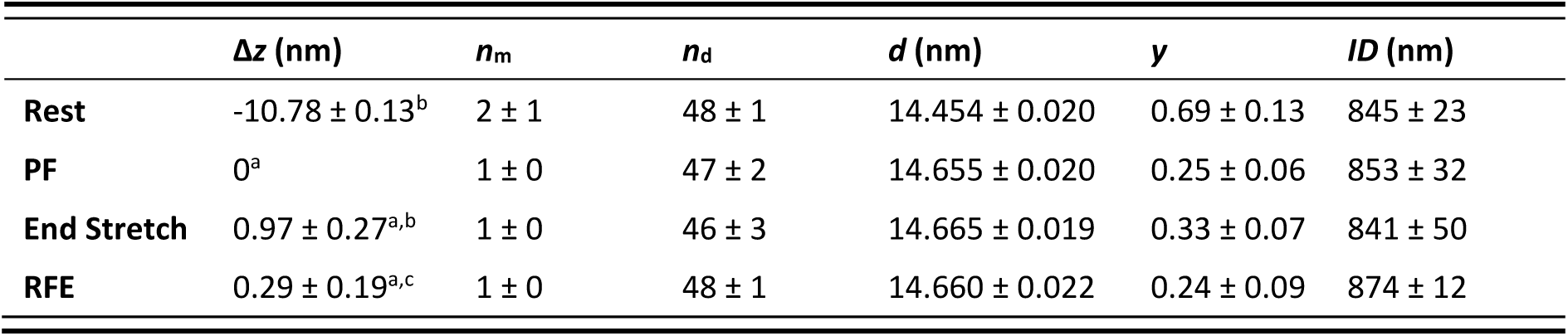
Best-fit model parameters in the interpretation of the axial profile of the M3 reflection. The rows denote the time periods in the protocol defined in Fig. 1a. The change in the axial centre of mass (Δ*z*) is shown relative to the half-bare zone value at peak force which was 90.96 ± 0.18 nm; the number of medial (*n*_m_) and distal (*n*_d_) layers of myosin motors marking the ends of the region of ordered motors; axial periodicity of myosin motors (*d*); scaling factor (*y*); interference distance (*ID*). Mean ± S.D. for n=5 muscles. Superscript characters denote significant Tukey’s post-hoc differences with respect to each key phase for Δ*z*; ^a^P<0.05 when compared to rest, ^b^P<0.05 when compared to peak force, ^c^P<0.05 when compared to end stretch.

| | $\Delta z$ (nm) | $n_m$ | $n_d$ | $d$ (nm) | $y$ | $ID$ (nm) |
| --- | --- | --- | --- | --- | --- | --- |
| Rest | $-10.78 \pm 0.13^b$ | $2 \pm 1$ | $48 \pm 1$ | $14.454 \pm 0.020$ | $0.69 \pm 0.13$ | $845 \pm 23$ |
| PF | $0^a$ | $1 \pm 0$ | $47 \pm 2$ | $14.655 \pm 0.020$ | $0.25 \pm 0.06$ | $853 \pm 32$ |
| End Stretch | $0.97 \pm 0.27^{a,b}$ | $1 \pm 0$ | $46 \pm 3$ | $14.665 \pm 0.019$ | $0.33 \pm 0.07$ | $841 \pm 50$ |
| RFE | $0.29 \pm 0.19^{a,c}$ | $1 \pm 0$ | $48 \pm 1$ | $14.660 \pm 0.022$ | $0.24 \pm 0.09$ | $874 \pm 12$ |

Further support for that conclusion was provided by the spacings of the myosin-based M3, M6 and M11 meridional reflections, *S*_M3_ (Fig. 3c), *S*_M6_ (Fig. 3d) and *S*_M11_ respectively (Fig. 3e). All these spacings increased by about 1.5% during fixed-end force development, signalling the longer axial periodicity of the activated state of the thick filament, but did not change significantly during the stretch. *S*_M11_, which is also associated with the ca 4 nm domain repeat of titin in the C-zone of the thick filament, did seem to increase transiently during the rapid force decline after the stretch (Fig. 3e).

### Thin filament regulation

The regulatory state of the thin filaments in muscle can be monitored using the intensity of the second actin-based X-ray layer line (*I*_AL2_)^24,25^. *I*_AL2_ increases during fixed-end force development, signalling the azimuthal movement of tropomyosin towards the centre of the groove between the two long-pitched strands of the thin filament to uncover the myosin binding sites on actin^26^. The signal-to-noise ratio of the *I*_AL2_ measurement was low (Fig. 4a), because of the axial dispersion of the AL2 layer line and the high background scattering in mouse EDL muscles. The mean value of *I*_AL2_ at the end of the stretch was 65% larger than that at peak force during fixed-end contraction, but this change was not significant at the p<0.05 level (Table S1).

**Fig. 4.**
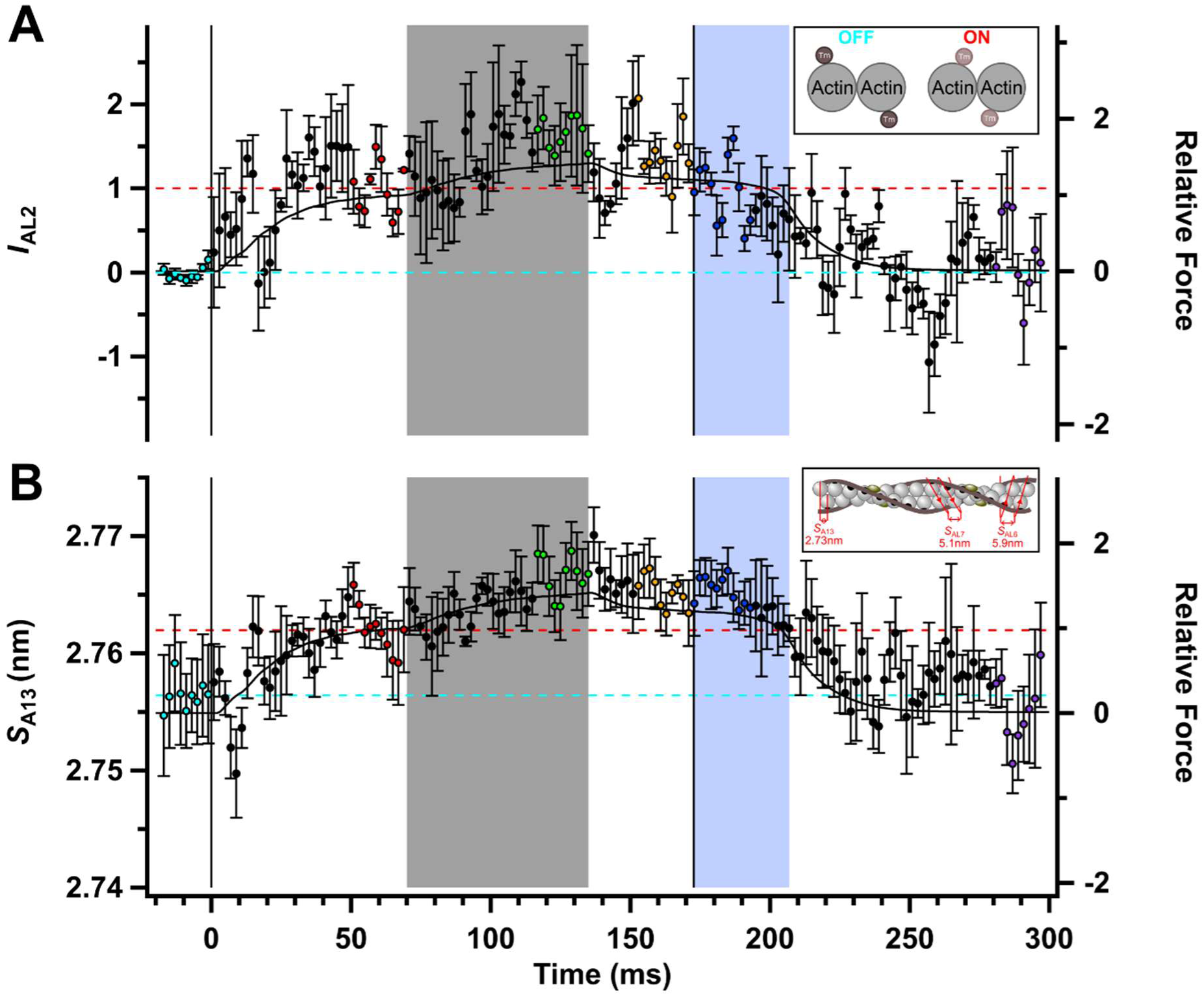
X-ray signals associated with thin filament activation. **a** Intensity of the second actin layer line (*I*_AL2_). *Inset*: Schematic transverse section through the thin filament indicating azimuthal positions of tropomyosin (Tm) at rest (cyan; dark brown circles) and in active contraction (red; light brown circles). **b** Spacing of the thirteenth meridional actin-based reflection (*S*_A13_), calculated from the spacings of the sixth and seventh actin layer lines. *Inset*: Longitudinal thin filament schematic with ca 2.73 nm axial repeat of actin monomers associated with the A13 reflection, and genetic actin helices associated with the AL6 (5.9 nm) and AL7 (5.1 nm) reflections. Tropomyosin (dark brown) blocks myosin binding sites on actin (black) when Ca^2+^ is not bound to troponin. Relative force (*T*/*T*_0_) is shown as the continuous black line. Colour coding of data points as in Fig. 1. Vertical lines denote the start and end of electrical stimulation; horizontal dashed lines denote the average resting value (cyan) and pre-stretch peak force value (red) of each parameter. Mean ± S.E.M. for n = 3 muscles in **a** and n = 5 muscles in **b**.

The axial spacing of actin monomers in the thin filament, called *S*_A13_ in reference to a model of the thin filament in which there are 13 actin monomers in each 37 nm helical repeat, has also been proposed to signal thin filament activation^27,28^. *S*_A13_ was larger at the end of the stretch than at peak force during fixed-end contraction (Fig. 4b; Table S1), and this significant (p<0.05) increase was maintained during residual force enhancement. However the mean increase is only 0.003 nm, of which 0.002 nm is the expected contribution from thin filament compliance^27,29^.

### The fraction of myosin motors attached to thin filaments

The most direct and specific X-ray signal for actin-attached force-bearing myosin motors is the intensity of the first actin-based layer line (*I*_AL1_)^30^, although the measurement of *I*_AL1_ is challenging because of high background and partial overlap with the first myosin layer line (Fig. S1c). *I*_AL1_ increased more than two-fold during fixed-end force-development (Fig. 5a; Table S1) as reported previously^22^. Its mean value decreased by 27% during the stretch and partially recovered afterwards, although these changes were not significant at the p<0.05 level (Table S1). Two other actin-based layer lines, AL6 and AL7, became more intense during fixed-end force development (Fig. 5b,c, Table S1). These layer lines are more clearly separated from myosin-based layer lines (Fig. S1e), and there were no significant changes in *I*_AL6_ and *I*_AL7_ during the stretch.

**Fig. 5.**
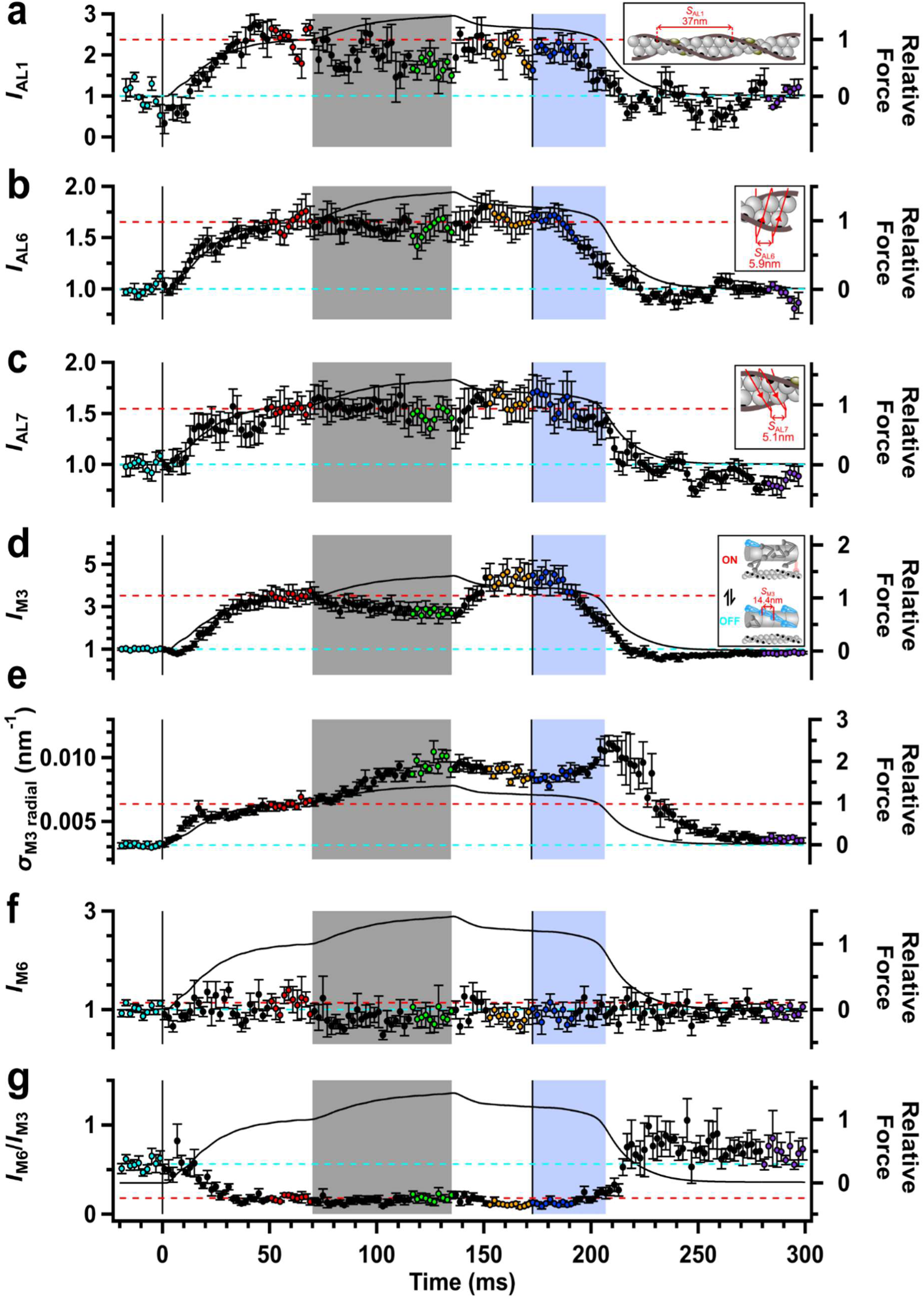
X-ray signals of myosin motor conformation and actin attachment. **a** Intensity of the first actin layer line (*I*_AL1_). *Inset*: Thin filament schematic showing the ca 37 nm helical periodicity associated with the AL1 reflection. **b** Intensity of the sixth actin layer line (*I*_AL6_) corresponding to the genetic helix with a periodicity of 5.9 nm as shown in the inset. **c** Intensity of the seventh actin layer (*I*_AL7_) corresponding to the genetic helix with a periodicity of 5.1 nm as shown in the inset. **d** Intensity of the third myosin-based meridional reflection (*I*_M3_) *Inset*: Schematic of overlapping thin and thick filaments with myosin motors in their resting (blue; axial periodicity 14.45 nm), actin-attached force-generating (light red), and disordered (dark grey) conformations. **e** Radial width of the M3 reflection (σ_M3 radial_). **f** Intensity of the sixth myosin-based meridional reflection (*I*_M6_). **g** Ratio of the intensities of the M6 and M3 reflections (*I*_M6_/*I*M3). Relative force (*T*/*T*_0_) is shown as the continuous black line. Colour coding of data points as in Fig. 1. Vertical lines denote the start and end of electrical stimulation; horizontal dashed lines denote the average resting value (cyan) and pre-stretch peak force value (red) of each parameter. Mean ± S.E.M. for n = 5 muscles for all data.

### The conformation of the myosin motors

The intensity of the M3 reflection (*I*_M3_; Fig. 5d), associated with the ca 14.5 nm axial periodicity of the myosin motors along the thick filaments, is the most widely used X-ray signal for changes in the conformation of the myosin motors^8,31^, although it is also sensitive to axial misalignment between neighbouring thick filaments, which can be quantified by the radial width of the reflection (σ_M3 radial_) (Fig. 5e). The decrease in width-corrected *I*_M3_ during the first 10 ms of fixed-end force development signals the release of motors from the folded OFF state on the surface of the thick filament; the later increase signals their binding to thin filaments in a conformation roughly perpendicular to the filament axis^22,32^. *I*_M3_ decreased during the stretch (Fig. 5d, green) as shown previously in whole muscles^33^ and single fibres^34^ from frog muscle, but this change was more than reversed after the stretch (orange). σ_M3 radial_ roughly doubled during fixed-end force development (Fig. 5e, red) and increased again by about the same amount during the stretch (green), indicating further axial misalignment between neighbouring thick filaments, which was only partly reversed after the stretch (orange).

Interpretation of these changes in *I*_M3_ is constrained by the splitting of the M3 reflection into three sub-peaks (Fig. 6a) as a result of X-ray interference between the two arrays of motors in each thick filament (Fig. S3). The relative intensity of the sub-peaks measures the movement of the centre of mass of the motors towards the midpoint of the thick filament^35^. During fixed-end force development, the fractional intensity of the low-angle (LA) peak of the M3 reflection (*L*_M3_; Fig. 6b) increased after a short lag phase, while that of the medium-angle MA peak decreased (*M*_M3_; Fig. 6c). The high-angle (HA) peak became very weak, again after a short delay (*H*_M3_; Fig. 6d). These changes can be explained by net axial movement of the centre of mass of the myosin motors by 11 nm away from the midpoint of the thick filaments (the M band of the sarcomere) during fixed-end force development (Fig. S3)^22,32^.

**Fig. 6.**
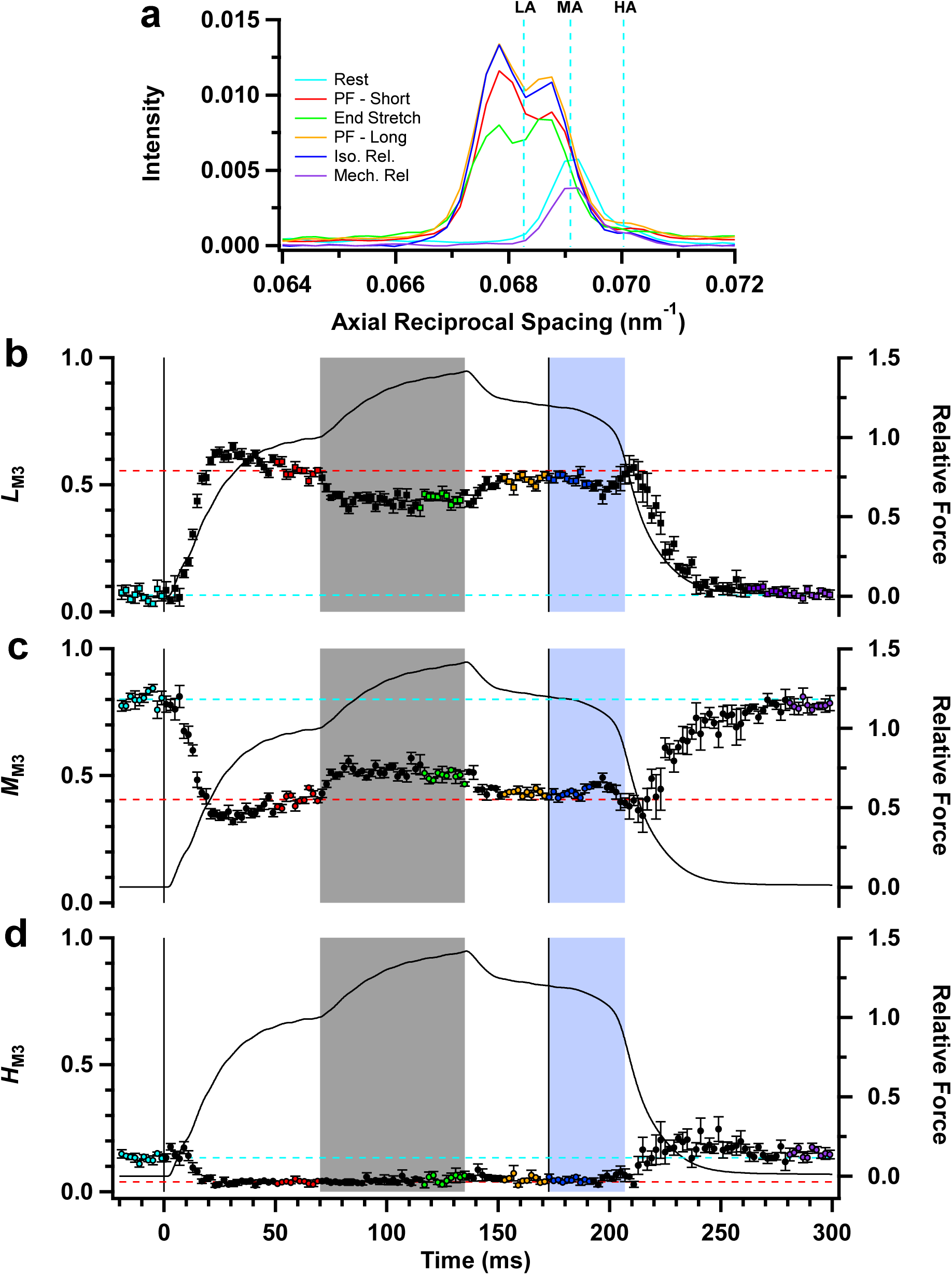
The fine structure of the M3 reflection. **a** Axial intensity distribution of the third myosin-based meridional reflection (M3), showing lower-angle (LA), mid-angle (MA) and higher-angle (HA) sub-peaks (vertical dashed cyan lines at rest). **b** Fractional intensity of the LA peak (*L*_M3_). **c** Fractional intensity of the MA peak (*M*_M3_). **d** Fractional intensity of the HA peak (*H*_M3_). Relative force (*T*/*T*_0_) is shown as the continuous black line. Colour coding of data points as in Fig. 1. Vertical solid lines denote the start and end of electrical stimulation; horizontal dashed lines denote the average resting value (cyan) and pre-stretch peak force value (red) of each parameter. Mean ± S.E.M. for n = 5 muscles for all data.

*L*_M3_ decreased during the stretch and *M*_M3_ increased, signalling that the centre of mass of the motors moved away from the midpoint of the thick filament. This movement was complete within the first 10 ms of stretch, when the relative sliding between thick and thin filaments is only about 7 nm, suggesting that myosin motor conformations had already reached a new steady state. Similarly, these reciprocal changes in *L*_M3_ and *M*_M3_ were reversed within 10 ms of the end of the stretch. The changes in *L*_M3_ and *M*_M3_ associated with the stretch were reproduced in a simple bipolar model of the thick filament (Fig. S3)^22^ by an axial movement (Δ*z*) of the centre of mass of the motors by 0.97 ± 0.27 nm away from the filament midpoint during the stretch (Table 1). This movement was largely recovered after the stretch, and there was no significant difference between the values of Δ*z* during residual force enhancement and that at peak force before the stretch. There was also no change in the interference distance, which was not significantly different from that expected for diffraction by all 49 motors in each half-filament in each phase of the protocol (Table 1). The axial width of the interference sub-peaks of the M3 reflection (σ_M3 axial_; Fig. S4) corresponds to a dispersion of the M3 periodicity of about 0.07 nm in resting muscle, increasing to about 0.08 nm during fixed-end force development, with no further significant change during or after the stretch.

The intensity of the M6 reflection (*I*_M6_; Fig. 5f), which is nominally the second order of the M3, did not change during fixed-end force development but decreased during the stretch. The ratio of the intensities of the M6 and M3 reflections (*I*_M6_/*I*_M3_; Fig. 5g) decreased during fixed-end force development, which may signal the lower axial dispersion of actin-attached myosin motors compared with that of the OFF motors^36^ but did not change significantly during the stretch. *I*_M6_/*I*_M3_ decreased during residual force enhancement after stretch (Fig. 1g), indicating a decrease in the conformational dispersion of the myosin motors.

### The time course of relaxation and its structural basis

Mechanical relaxation is slower after tetani containing a stretch (Fig. 7). The isometric phase of relaxation is prolonged, and the transition to the exponential phase is delayed (Fig. 7a, blue) compared with that in fixed-end tetani (black)^22^, or tetani containing a period of unloaded shortening (red)^23^. That transition, which corresponds to an inflection point in the force trace (Fig. 7b), is correlated with the later appearance of two sarcomere populations (Fig. 7c). Relaxation started from a slightly longer sarcomere length (Fig. 7c) and smaller lattice spacing (Fig. 7d) in tetani containing a stretch, so one of these factors may be responsible for the delayed relaxation. The slower relaxation is also associated with slower initial decreases in *I*_1,1_/*I*_1,0_ (Fig. 7e), *I*_M3_ (Fig. 7f) and *S*_M3_ (Fig. 7g) after the last stimulus, indicating that myosin motors detach more slowly from thin filaments after a stretch. The triphasic changes in σ_M3 radial_ (Fig. 7h) and *L*_M3_ (Fig. 7i) after the last stimulus are also delayed after a stretch, which may signal strain-dependent detachment kinetics within the population of actin-attached motors.

**Fig. 7.**
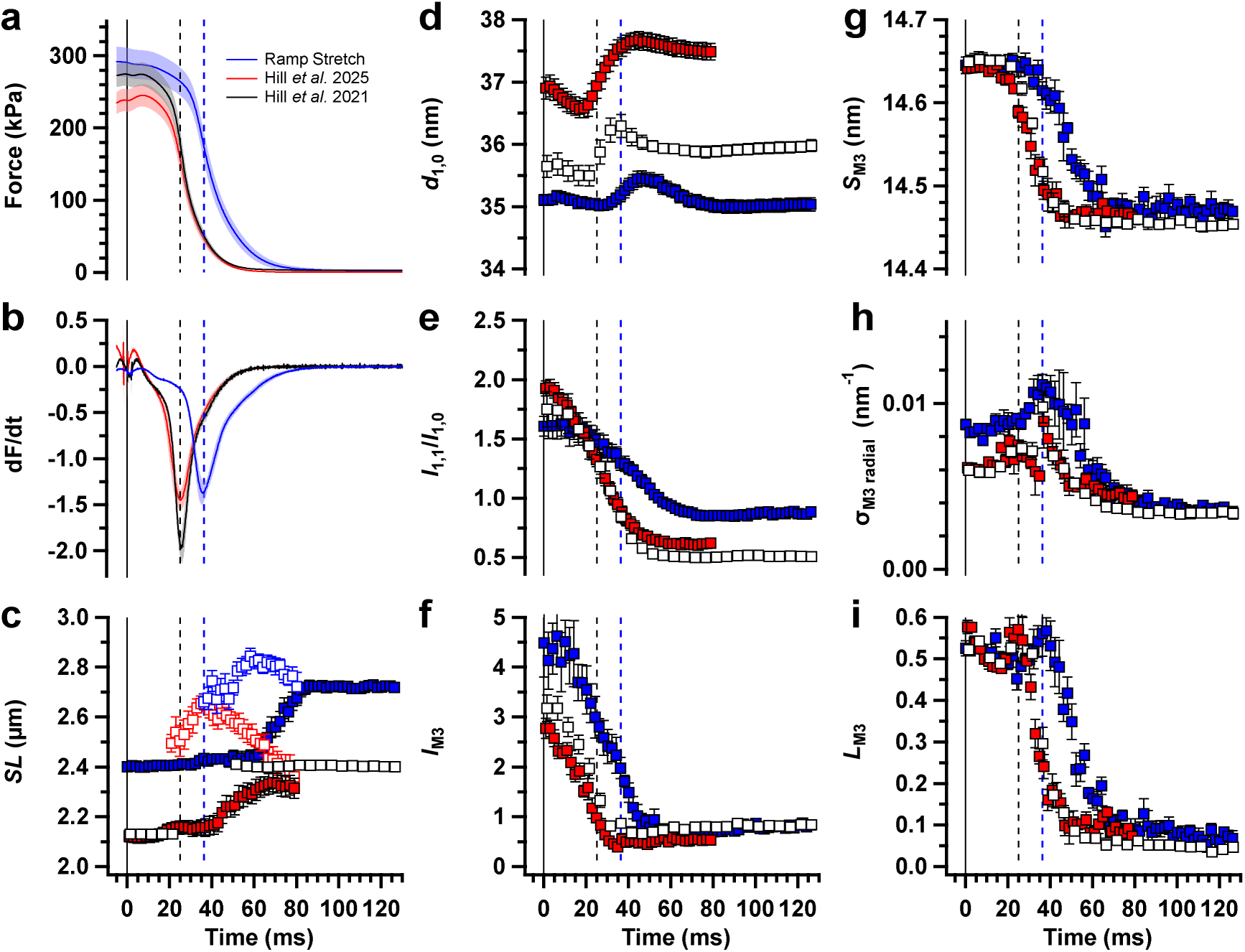
Comparison of force and X-ray signals during relaxation in tetani with different imposed length changes. **a** Relative force (*T*/*T*_0_) with respect to the last electrical stimulus (vertical black line; t = 0 ms) following a tetanus containing a ramp stretch (blue; this work), following a fixed-end tetanus (black, Hill *et al*.^22^) and following a tetanus containing unloaded shortening (red, Hill *et al*.^23^), all in mouse EDL muscle at 27°C. **b** First differential of the force data in **a** (dF/dt). **c** Sarcomere length (*SL*). **d** Lattice spacing (*d*_1,0_). **e** Equatorial intensity ratio (*I*_1,1_/*I*_1,0_). **f** Intensity of the third myosin-based meridional reflection (*I*_M3_). **g** Spacing of the third myosin-based meridional reflection (*S*_M3_). **h** Radial width of the M3 reflection (σ_M3 radial_) **i** Fractional intensity of the LA peak of the M3 reflection (*L*_M3_). For **c**-**i**, filled blue squares, this work; open white squares, Hill *et al*.^22^; filled red squares, Hill *et al*.^23^. Vertical dashed lines denote the time of minimum dF/dt in the ramp stretch protocol (blue) and the other protocols (black), respectively. For **d** and **g**, the spacings published in Hill et al.^22^ and Hill *et al*.^23^ were recalculated using the beamline calibrations. Data are presented as mean ± S.E.M. for 5-7 muscles for each protocol.

## Discussion

The experiments described above were designed to elucidate the molecular structural basis of the braking action of muscle. To do so, we used time-resolved X-ray diffraction to reveal the changes in the conformation of the myosin motors, the regulatory states of the thin and thick filaments, and the organisation of the filament lattice. We applied a constant velocity stretch to intact EDL muscles of the mouse during electrical stimulation in the conditions we had previously used to characterise the molecular structural changes in fixed-end and shortening contractions^22,23,36^. The stretch was applied at a sarcomere length close to the optimal for fixed-end force production, and its amplitude and speed were limited to avoid the sarcomere inhomogeneity that can contribute to the stretch response in the less physiological conditions of long sarcomere length and large or fast stretches.

We confirmed the homogeneous response of the muscle sarcomeres to stretch in the present experiments by ultra-small angle X-ray diffraction. In contrast with diffraction of visible light, the lower scattering probability of the sarcomeres at X-ray wavelengths allows the full distribution of sarcomere lengths throughout the cross-section of an intact muscle to be determined. The stretch produced an average sarcomere length increase of 0.08 µm, corresponding to filament sliding of about 40 nm in each half-sarcomere, and these changes were reproducible along and between muscles. Sarcomere length dispersion increased by only 0.02 µm during the stretch, confirming the conclusion of earlier studies with single muscle fibres^11,16^ that sarcomere inhomogeneity makes a negligible contribution to the force response to slow stretch near optimal muscle length.

In the absence of sarcomere inhomogeneity, the braking action of muscle in response to stretch must be an intrinsic property of the sarcomere. In the widely accepted cycling cross-bridge paradigm of muscle contraction^2^ and its later elaboration to include a powerstroke in the actin-attached myosin motor^3,37^, the increased force during stretch might be due to either an increase in the number of myosin motors bound to actin or an increase in the force per motor. However, the mechanical and energetics studies described in the Introduction^13,15^ show that the braking action of the sarcomere, and in particular the energy stored during the stretch and the residual force enhancement after it, cannot be explained solely by changes in the myosin motors. We therefore addressed two complementary questions in the present study: (i) how much of the force enhancement during and after stretch can be explained by an increase in the number of actin-attached motors or the force per attached motor?; and (ii) how does the stretch alter the structural organisation of the filament lattice and the sarcomeric cytoskeleton?

The most direct X-ray signal for the fraction of myosin motors attached to actin is the intensity of the first actin-based layer line (*I*AL1), which is almost linearly related to that fraction^30^. *I*_AL1_ decreased by about 30% during the stretch, and although this decrease was not significant at the p<0.05 level, it is consistent with the 38% decrease reported for slow stretch of demembranated fibres from rabbit psoas muscle at 35°C^38^. Such a decrease in *I*_AL1_ is not consistent with the hypothesis that the extra force during stretch is due to an increase in the fraction of myosin motors attached to thin filaments, and several other X-ray signals confirm this conclusion. The intensities of the sixth and seventh actin-based layer lines, which are also sensitive to the fraction of attached motors, did not change during stretch. The intensity of the 1,0 equatorial reflection (*I*_1,0_) did not change during stretch and that of the 1,1 reflection (*I*_1,1_) decreased, as reported previously for frog muscle^20^, in contrast with the decrease in *I*1,0 and increase in *I*_1,1_ that would be expected for increased motor attachment.

The conclusion from multiple X-ray signals that the fraction of myosin motors attached to thin filaments does not increase during stretch and may decrease slightly is inconsistent with previous interpretations of the roughly 20% increase in sarcomere stiffness^11,35,39^. If the observed stiffness increase were solely due to a greater fraction of myosin motors attached to thin filaments, the substantial compliance of filaments themselves means that the fraction of attached motors would have to increase by about 80% during the stretch^35^, which is clearly inconsistent with the X-ray results. Those results therefore suggest that the increase in sarcomere stiffness during stretch is associated with distortion of the filament lattice and sarcomeric cytoskeleton^15,17,40^ rather than an increase in the number of actin-bound myosin motors.

The regulatory state of the thick filaments does not change during the stretch, as shown by multiple X-ray signals for thick filament activation, including the intensities of the myosin-based layer lines associated with the helical arrangement of the myosin motors in the OFF state of resting muscle and the axial periodicity of the thick filament reported by the spacings of the M3, M6 and M11 reflections. This result is in marked contrast to the *in*activation of the thick filaments during shortening^23^. Thick filaments are activated through a mechano-sensing mechanism^41^, but the present results show that mechanism is limited to forces less than that in a fixed-end contraction. Thick filaments are already fully ON during fixed-end force generation, and the 40% increase in force during the stretch does not release additional motors for thin filament binding.

We also attempted to determine the regulatory state of the thin filaments during the stretch using the intensity of the second actin-based layer line, which is sensitive to the azimuthal movement of tropomyosin that controls the availability of myosin binding sites on actin. The mean value of *I*_AL2_ increased during the stretch, but the signal-to-noise was so low that we could not distinguish between two alternative hypotheses: that *I*_AL2_ is unaffected by the stretch or that it increases by 40% in proportion to the force. Further experiments will be required to determine the effect of stretch on the level of thin filament activation.

If the number of myosin motors bound to thin filaments does not increase during stretch, as argued above, the increase in force must either be due to an increased force per motor, distortion of the cytoskeleton, or both. We addressed the first possibility using X-ray interference to determine the conformation of the actin-attached myosin motors during the stretch, which is signalled by the fractional amplitude of the low-angle sub-peak of the M3 reflection (*L*_M3_)^7,8,22^. *L*_M3_ decreased during the first 10 ms of the stretch, corresponding to the first 7 nm of filament sliding, but was constant during the remainder of the stretch, indicating a constant average motor conformation as filaments continue to slide and actin-attached motors must be repeatedly strained, detach from actin, and reattach to different actin monomers.

This steady-state change in *L*_M3_ produced by stretch corresponds to the average centre of mass of the myosin motors being about 1 nm further away from the midpoint of the thick filament than during fixed-end contraction (Table 1). Exactly the same centre-of-mass displacement was estimated previously from the change in *L*_M3_ during slow stretch of demembranated muscle fibres from rabbit psoas muscle at 37°C^38^. Since about half of the myosin motors that contribute to the M3 X-ray reflection are not displaced in response to filament sliding^7,9^, the centre of mass of the actin-bound motors must move by about twice that amount. In the widely accepted structural model for the powerstroke^6^, the relative masses of the motor components mean that a 2-nm displacement of the centre of mass requires rotation of the light chain domain with respect to the actin-bound catalytic domain to displace its distal end about 2.4 nm. Tilting of the light-chain domain has both elastic and powerstroke components^42^, and the contribution of the elastic component to the first 10ms of the stretch response, during which the steady state is established, can be estimated from the observed force increase and the motor compliance^43^ as about 0.6 nm. The remaining 1.8 nm of the observed conformational change in the actin-attached motor is therefore likely to be associated with partial reversal of the powerstroke by the stretch. We conclude that the fast component of the force increase during the first 10ms of stretch could be due to the increased strain of actin-attached myosin motors.

This steady-state conformational change in the myosin motors during slow stretch is smaller than that produced by much faster (roughly 100 µs) stretches used in previous X-ray interference experiments in intact fibres from frog muscle^44^. The observed changes in the interference fine structure of the M3 reflection during and immediately after those large fast stretches provided evidence for attachment of the second motor domain of each myosin molecule to the thin filament. That conclusion was based on the kinetic separation of the elastic and powerstroke components made feasible by the step stretch, which is not possible with the smaller and much slower stretches used here. Although we cannot exclude the possibility that the second motor domain attaches transiently during the fast component of the response to a slow stretch, the present results show that any such recruited motors have detached in the steady-state response to the much slower stretches used here. Both speeds of stretch are likely to be physiologically relevant, and the attachment of the second motor domain may be a specific adaption for fast stretch, in which speed of the response is paramount, compared to slow stretch, when metabolic efficiency is likely to be the dominant driver.

The slow component of the force increase during slow stretch, after the steady state of motor strain has been established, cannot be due to the myosin motors. Three X-ray signals suggest that this component is due to perturbation of the lattice of thin and thick filaments and their lateral connections in the Z-disk and M-band of the sarcomere (Fig. 8). First, the selective decrease in *I*_1,1_ during the stretch suggests that thin filaments are displaced from the trigonal positions of the hexagonal filament lattice (Fig. 8c)^20^. Those displacements might be related to the intrinsic instability of the thin filament position pulled in three different directions by the neighbouring myosin motors, or to increased binding of thin filaments to titin in the I-band^40^. Second, the inter-filament spacing and filament lattice volume decrease during stretch (dashed lines in Fig. 8c, denoting the larger lattice in Fig. 8a), which implies lateral compression of the M-band. Third, the radial width of the M3 reflection increases beyond that during fixed-end contraction, indicating larger axial displacements between neighbouring thick filaments (Fig. 8b,d), which would distort the transverse links between thick filaments in the M band. The radial and axial forces associated with these three effects could potentially be connected^45^ and contribute to elastic energy storage during stretch and the increase in axial stiffness. Moreover, all three effects continue during the slow phase of the response to stretch, and are incompletely reversed after stretch, so they could also contribute to residual force enhancement after stretch.

**Fig. 8.**
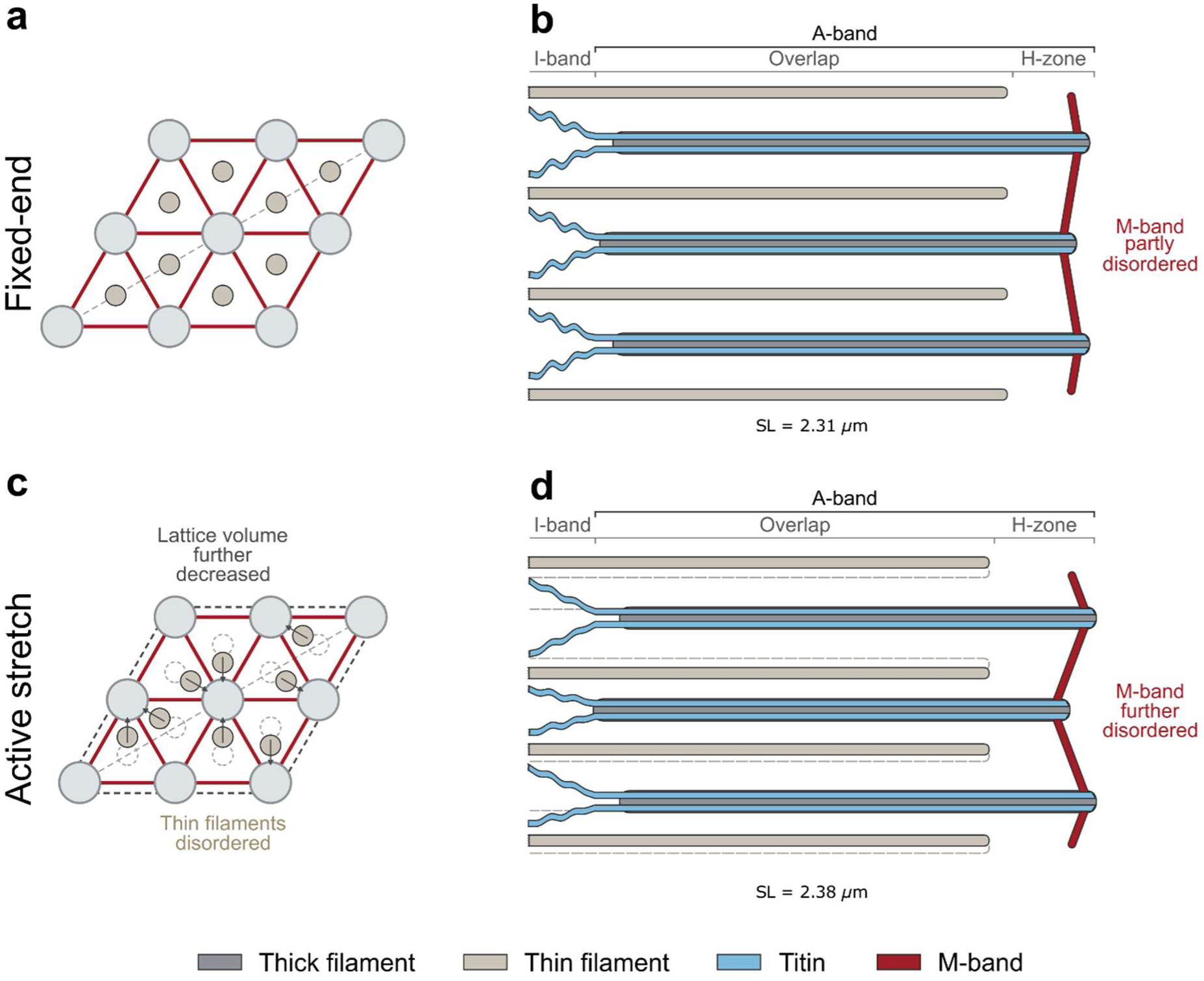
Perturbation of the lattice of thick and thin filaments during stretch. **a** Cross-section of the hexagonal lattice of thick (light grey) and thin (gold) filaments in the overlap region of the A-band during fixed-end contraction. Links between thick filaments at the M-band are in red. The ideal hexagonal lattice is drawn for simplicity, although there would be random displacements from these positions in practice. **b** Longitudinal view of the thick filaments (grey), thin filaments (gold) and titin (blue) in a half-sarcomere during fixed-end contraction. The actin-and titin-containing I-band, the filament overlap region in the A-band, and the H-zone are labelled. Sarcomere length (*SL*) in fixed-end contraction is already shorter than at rest, and small axial misalignments between neighbouring thick filaments are associated with some disorder of the M-band (red). **c** Cross-sectional view of the A-band during active stretch. Lattice volume decreases with respect to the fixed-end contraction (dashed dark grey rhombus), and actin filaments are displaced from their trigonal positions fixed-end contraction (dashed gold circles) towards the thick filaments, denoted by arrows. **d** Longitudinal view of part of the half sarcomere during active stretch. *SL* has increased and lattice volume has decreased with respect to the fixed-end contraction (horizontal dashed grey lines). The M-band region is more distorted and the axial misalignment between neighbouring thick filaments has increased. Thin filaments become displaced from their original positions (dashed gold lines).

In contrast with the rapid recovery of the stretch-induced increase in motor strain after the end of the stretch, all three of these X-ray signals of filament lattice distortion do not recover after the end of the stretch. We therefore propose that filament lattice distortion is responsible for the residual force enhancement after stretch as well as for the slow component of the force increase during stretch. This would also explain why residual force enhancement can be maintained for several seconds after the end of the stretch^16^, much longer than the ATPase cycle of the myosin motors. However, one striking change in the actin-attached myosin motors is retained after the stretch, signalled by the maintained increase in *I*_M3_ to a level greater than that at the peak of force during fixed-end contraction before the stretch. This is accompanied by a significant decrease in the dispersion of the myosin motors signalled by an increase in *I*_M6_/*I*_M3_ (Table 1), despite the fact that the average conformation of the motors, signalled by *L*_M3_, is the same. These results suggest that the cycles of motor detachment and reattachment during stretch result in a narrowing of the spread of motor conformations, which may also contribute to the residual force enhancement.

The delayed mechanical relaxation from the residual force-enhanced state suggests a slower rate of detachment of myosin motors from thin filaments. Since the detachment rate is likely to be strain-dependent, this effect might be a consequence of the lower conformational dispersion of the myosin motors after the stretch, but it could also be related to another effect of filament lattice distortion.

## Methods

### Muscle Preparation

Male C57BL/6 J mice aged 4 to 8 weeks were housed and maintained at the former ID17 Biomedical Facility at the ESRF in 12:12 hour light:dark cycles at 50% relative humidity for one week prior to experiments. Animals were sacrificed via cervical dislocation followed by a confirmation method of permanent cessation of circulation via severing the femoral artery in accordance with the European Union regulation (directive 2010/63). Intact mouse EDL muscles were dissected and mounted, via hooks, vertically in a custom 3D-printed trough at beamline ID02 of the ESRF (Grenoble, France)^46^, attached to a dual-mode force/length transducer (300C-LR, Aurora Scientific, Aurora, Canada) and continuously perfused with Krebs–Henseleit solution equilibrated with carbogen at 28°C. Electrical stimuli were provided by a high-power biphasic stimulator (701C, Aurora Scientific) via parallel platinum electrodes. The stimulus voltage was set to 1.5 times the required amount to elicit the maximum twitch force response, and optimal muscle length (*L*_0_) was set to produce maximum force in response to a 100 ms train of stimuli at 110 Hz. Muscle wet weight and *L*_0_ were 10.7 ± 0.6 mg and 13.8 ± 0.1 mm, respectively. The cross-sectional area was 1.5 ± 0.1 mm^2^. The maximal isometric force of the muscles before exposure to radiation was 301.9 ± 12.9 kPa.

### Ultra-small-angle and Small-angle X-ray Diffraction Data Collection

Ultra-small-angle and small-angle X-ray diffraction patterns were recorded using an Eiger 2 4M detector (Dectris Inc., Baden, Switzerland) consisting of a 2 by 4 array of detector tiles with small gaps between them, and normalised for the detector response, incident intensity, and diffraction geometry^46^. The sample-to-detector distance was initially set to 31 m to collect ultra-small-angle X-ray diffraction patterns for the measurement of sarcomere length (*SL*), then to either 3.2 or 2.0 m to record small-angle X-ray reflections associated with filament periodicities in the range 60 - 3.5 nm^23^.

Muscles were aligned in the X-ray beam using a sample-to-detector distance of 31 m using 1 ms exposures with a 50 μm rhodium attenuator. Following alignment, the rhodium attenuator was replaced with a 20 μm lead attenuator.

Muscles were held at *L*_0_ and stimulated for 172 ms at 110 Hz. After the first 70 ms of electrical stimulation under fixed-end conditions, the muscle was stretched by 2.5% of *L*_0_ in 65 ms whilst stimulation continued, then held at the new length for 36 ms until the final electrical stimulus was delivered at 173 ms. The stimulus and length change protocol is shown in Fig. 1a. X-ray diffraction data were acquired in 160 time frames, each with 1.8 ms integration and 0.2 ms latency time. To minimise radiation damage, X-ray exposure was limited by a fast shutter, and muscles were moved vertically and horizontally between successive X-ray exposures. X-ray data were averaged from 15 to 35 contractions per muscle. Records in which force had declined more than 15% from the first stimulus were excluded from further analysis.

Force, stimulus, muscle length, and X-ray acquisition timing were sampled and analysed using custom-made software written in LabVIEW (National Instruments).

### X-ray Diffraction Data Analysis

Normalised data were analysed using SAXSutilities2, SAXS package (P. Boesecke, ESRF, Grenoble, France), Fit2D (A. Hammersley, ESRF, Grenoble, France), ImageJ (National Institute of Health, Bethesda, MD)^47^ and Igor Pro (v8 and v10, WaveMetrics, Inc., Portland, OR). Small-angle X-ray diffraction patterns containing collagen-based reflections (∼2 to 6 per muscle), indicating the presence of tendons in the X-ray beam, were excluded from the analysis.

The instrument background was subtracted from each normalised pattern, and useable records from the same muscle were averaged. Patterns were corrected for any tilting of the muscle with respect to the detector pixels using a custom-written ImageJ macro.

### Analysis of Ultra-Small-Angle X-Ray Diffraction Patterns

The sarcomeric X-ray reflections were recorded with a sample-to-detector distance of 31 m at three places along the long axis of each muscle. Diffraction data were integrated 0.56 µm^−1^ on either side of the meridional axis, and the residual background intensity was removed using an arc hull algorithm using the Baselines extension in Igor Pro, with depth and smoothing set to zero. The axial region 0.20-0.59 µm^−1^ containing the first-order sarcomere reflections was fitted with a single Gaussian function for time frames up to 207 ms (Fig. 1c), and a small correction was made for the beam size and effective point spread function of the detector. σ for sarcomere length (σ_SL_) was then calculated assuming that the fractional dispersion in sarcomere length is equal to that of the corrected intensity distribution on the detector. At later times during relaxation, two peaks could be resolved in this region, which were fitted with two Gaussians with different axial widths (Fig. 1b,c).

### Equatorial reflections

The equatorial intensity distribution was determined by axially integrating 0.0036 nm^−1^ on either side of the equatorial axis (perpendicular to the muscle axis), and the residual background intensity removed using an arc hull algorithm using the Baselines extension in Igor Pro, with depth set to zero and smoothing set to two. The intensities, radial positions, and widths of the 1,0, Z-disc, 1,1 and 2,0 reflections were determined by fitting four Gaussian peaks to the radial region 0.02 to 0.065 nm^−1^. Peak positions were constrained according to the hexagonal filament lattice geometry:

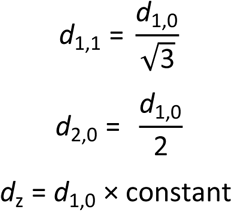

where *d*_h,k_ represents the lattice spacing associated with the corresponding reflection.

The volume of the filament lattice in each sarcomere (*V*) was calculated as:

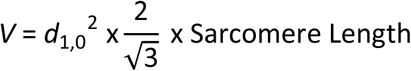

The radial widths of the 1,0 (σ_1,0_) and 1,1 (σ_1,1_) reflections were corrected for beam dimensions and effective point spread function of the detector and expressed in nm^−1^.

### Meridional reflections

Processed two-dimensional X-ray diffraction patterns were radially integrated 0.0075 nm^−1^ on either side of the meridional axis, and the M3, M6 and M11 meridional reflections fit in the axial regions 0.066-0.072 nm^−1^, 0.135-0.142 nm^−1^ and 0.246-0.260 nm^−1^, respectively. The residual background intensity was removed using an arc hull algorithm using the Baselines extension in Igor Pro, with depth set to zero and smoothing set to two. The interference components of the M3 and M6 reflections were determined by fitting multiple Gaussian peaks of equal axial width to the meridional intensity distributions. The cross-meridional widths of the M3 and M6 reflections were obtained from integrations of the above-defined axial regions and fit using a double Gaussian centred on the meridian with two separate widths, the smaller of which was assumed to be the best estimate of the radial width of the M3 or M6 reflection. For the M11 reflection, 1:2:1 smoothing of the integrated intensity as a function of time was performed to increase the signal-to-noise ratio. The M11 reflection was then fitted with a single Gaussian to obtain intensity and spacing. The intensities of the M3 and M6 reflections were calculated as the sum of their component peaks and multiplied by their cross-meridional widths to account for axial misalignment of thick filaments^48^. The M11 intensity could not be corrected by its cross-meridional width and is not reported.

### Layer line reflections

The intensities of the first myosin and first actin layer lines (ML1 and AL1) were calculated by integrating the radial region between 0.037 nm^−1^ and 0.064 nm^−1^ from the meridional axis using the 3.2 m data. Due to the partial overlap of ML1 and AL1, integrated intensities were obtained from the axial region 0.017-0.033 nm^−1^ for each muscle and separated by global Gaussian deconvolution of the time series data under the simplifying assumption that their spacings, *S*_ML1_ and *S*_AL1_ respectively, and axial widths do not change during contraction.

The intensity of the ML4 layer line was obtained by integrating radially in the reciprocal range 0.030-0.077 nm^−1^ parallel to the meridian. Due to the proximity of the ML4 reflection to a detector tile boundary in unmirrored data, this integration used only half of the pattern. Furthermore, an additional reflection, the meridional AM_+1_ reflection at ∼0.10 nm^−1^, partially overlaps the ML4 reflection on the high-angle side (Fig. S1d)^49^. The intensity of the ML4 reflection was therefore obtained by integrating the lower angle region of the reflection in the axial range 0.088-0.093 nm^−1^ (Fig. S1d, grey shading).

The intensity of the AL2 layer line was obtained by integrating the radial region 0.160-0.303 nm^−1^, then integrating axially in the region 0.0373-0.0729 nm^−1 50,51^. The second actin layer line was not detectable in resting muscle, so changes in its integrated intensity were normalised to the value at peak force before the stretch.

Intensity distribution profiles for the AL6 and AL7 reflections were obtained by integrating in the radial region 0.035-0.060 nm^−1 52^, with the lower limit chosen to minimise the contribution of the tails of the M7, M8 and M9 reflections (Fig. S1e). Due to the proximity of the AL6 reflection to a detector tile boundary, this integration was performed on half of the pattern. The integration for the weaker AL7 reflection was performed on fully mirrored patterns to maximise signal-to-noise ratio. The spacings and intensities of the AL6 and AL7 reflections were obtained by global Gaussian deconvolution of five Gaussian fits of independent axial spacing and width in the axial regions between 0.166-0.178 nm^−1^ for AL6 and 0.191-0.215 nm^−1^ for AL7, respectively^53^ assuming that the spacings of the M7 and M8 reflections and the axial width of all the reflections did not change during contraction.

The thirteenth-order actin-based meridional reflection (A13), which corresponds to the fundamental ∼2.7 nm axial repeat of actin monomers in the thin filament, was outside the detector in the present experiments, but its spacing was calculated from *S*_AL6_ and *S*_AL7_ using the equation^52^:

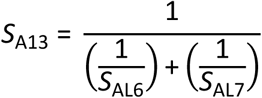

For all actin and myosin layer line reflections, the axially integrated 1D profiles were first 1:2:1 smoothed as a function of time to increase the signal-to-noise ratio of the weakly diffracting reflections. The residual background intensity was subtracted using an arc hull algorithm using the Baselines extension in Igor Pro, with depth set to zero and smoothing set to two for all layer line reflections.

### Correction of Integrated Intensities for changes in muscle mass in the X-ray beam

Small movements of the muscles with respect to the X-ray beam during contraction can alter the scattering mass in the beam and, consequently, the diffracted X-ray intensities. All reported X-ray intensity time-courses were corrected for this effect by normalising with respect to the background under the ML1 layer line at each frame. At the force plateau, the average correction factor was 1.04 ± 0.05 and it did not change during the stretch.

### Interpretation of the axial profile of the M3 reflection

Myosin filaments from skeletal muscle are centrosymmetric about their midpoint at the M-band of the sarcomere and contain two arrays of 49 myosin motors with an axial periodicity *d* of about 14.5 nm, corresponding to the spacing of the M3 X-ray reflection (*S*_M3_), with the first layer (layer 1) starting about 80 nm (the half bare zone, HBZ) from the filament midpoint. The centrosymmetric structure of the myosin filament generates an interference function that samples the X-ray reflection produced by a single array of motors to give finely spaced sub-peaks^35,54^, from which the length of each array of diffracting motors contributing to the M3 reflection and the distance between their centres – the interference distance *D* – can be determined. Each layer of myosin motors in the filament was represented as a point diffractor, which gives an excellent approximation to the profile of the M3 reflection calculated from the full axial mass profile of the motors^55^.

In the point diffractor model, HBZ represents the average centre of mass of the first layer of myosin motors from the filament midpoint, and the diffracted intensity distribution along the meridional axis can be calculated as^55^:

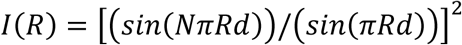

where *R* is the reciprocal space co-ordinate, *N* is the number of layers of motors in a contiguous array from the first medial layer, *n*_m_, to the last distal layer, *n*_d_, that contribute to the M3 reflection, while layers outside that region make zero contribution because their motors are isotropic, and *D* is the interference distance calculated as:

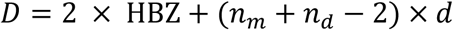

For fitting to the experimental M3 profiles, the calculated *I*(*R*) was convoluted with a Gaussian function with sigma 90 µm on the detector to mimic the effects of the axial dispersion of the M3 spacing (Fig. S4). The best fit for each muscle and time point was determined by a global search of HBZ, *d*, *n*_m_, *n*_d_ and an intensity scaling factor, *y*, by minimising *χ*^2^ calculated using the experimental standard deviations from the difference between the experimental mean and the model output for the more intense peaks at each time point, corresponding to a reciprocal space region ranging between ∼0.0662 and ∼0.0708 nm^−1^.

### Statistical Analyses

All statistical analyses were performed using Jamovi (The Jamovi Project, v2.7.37) and Microsoft Excel. Differences in force or X-ray data between the key time periods in the protocol shown in Fig. 1 were analysed using a repeated measures analysis of variance (ANOVA) with Tukey’s post hoc analysis for data where a main effect was observed (Table 1, Table S1). To determine whether non-parametric analyses were required, data were first checked for normality of distribution using Shapiro-Wilks test, skewness, kurtosis and sphericity using Mauchley’s W. Those which were not spherical used a Greenhouse-Geisser sphericity correction. Main effects and post-hoc analyses P-values are provided in Table S2. Data presented are mean ± standard error of the mean (S.E.M.) unless otherwise stated. Significance was set at P<0.05 for all analyses.

## Acknowledgements

We would like to thank the ID02 technical staff, Laurent Jacqmin, Peter Boesecke, and Michael Sztucki and the Biomedical Facility staff, Mélanie Jomard, Andrea Tramond and Michael Kirsch (European Synchrotron Radiation Facility), for their support during the beamtime; the European Synchrotron Radiation Facility for the award of synchrotron beamtime; Kawal Rhode and Zhouyang Xu (King’s College London) for their assistance with the 3D printing of the muscle trough. This work was funded by the Medical Research Council (MR/R01700X/1), awarded to M.I., and the European Synchrotron Radiation Facility. M.K. was supported by the Wellcome Trust (215482/Z/19/Z) award to M.I. A.A. was supported by King’s British Heart Foundation Centre of Research Excellence Award RE/18/2/34213 and FS/4yPhD/F/21/34154B. E.B. was supported by a British Heart Foundation Intermediate Basic Science Research Fellowship (FS/17/3/32604). LF was funded by a Sir Henry Dale Fellowship awarded by the Wellcome Trust and the Royal Society (210464/Z/18/Z).

## Author Contributions

Designed Research – CH, EB, LF, MI; Performed Research – CH, MK, AA, AF, TN, EB, LF, MI; Contributed new reagent/analytic tools – TN; Analysed data – CH, EB; Wrote the paper – CH, EB, LF, MI; Interpreted Results – CH, EB, LF, MI; Revised the Manuscript – CH, MK, AA, AF, TN, EB, LF, MI

---

**Fig. S1.**
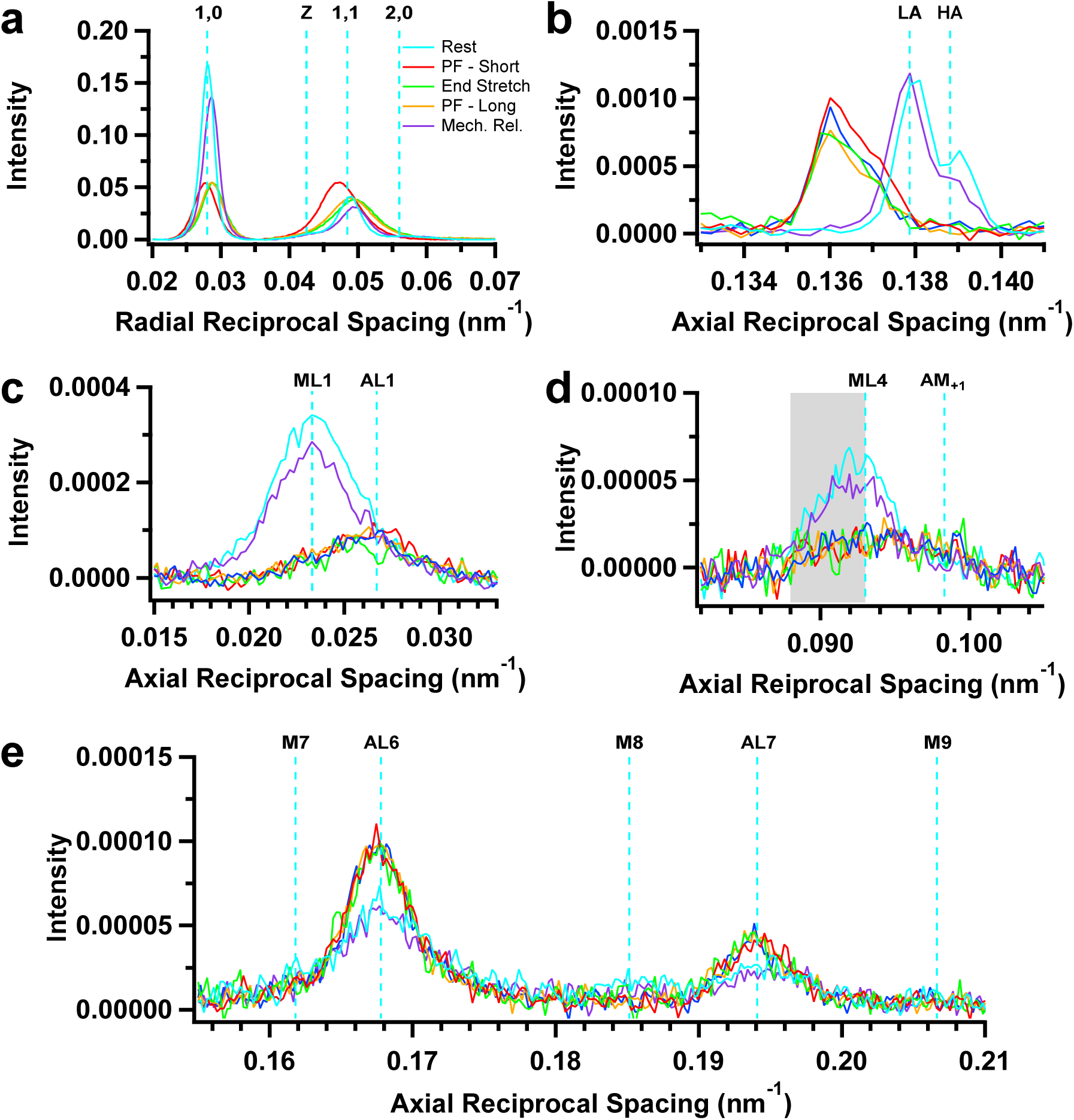
Intensity distributions of equatorial, meridional and layer line X-ray reflections. **a** Equatorial intensity distribution showing the 1,0, 1,1, 2,0 and Z-disc reflections. **b** Axial intensity distribution of the sixth meridional reflection (M6) showing the lower-angle (LA) and higher-angle (HA) sub-peaks denoted by vertical cyan lines at rest. **c** Axial intensity distributions in the region of the ML1 and AL1 layer lines. **d** Axial intensity distributions in the region of the ML4 and actomyosin (AM+1) layer lines. *I*_ML4_ was obtained from the grey-shaded region to avoid any contribution from AM_+1_. **e** Axial intensity distributions in the region of the AL6 and AL7 layer line reflections. The seventh, eighth and ninth meridional reflections (M7, M8 and M9, respectively) are indicated. Muscle-to-detector distance, 3.2 m; average from n=5 muscles.

**Fig. S2.**
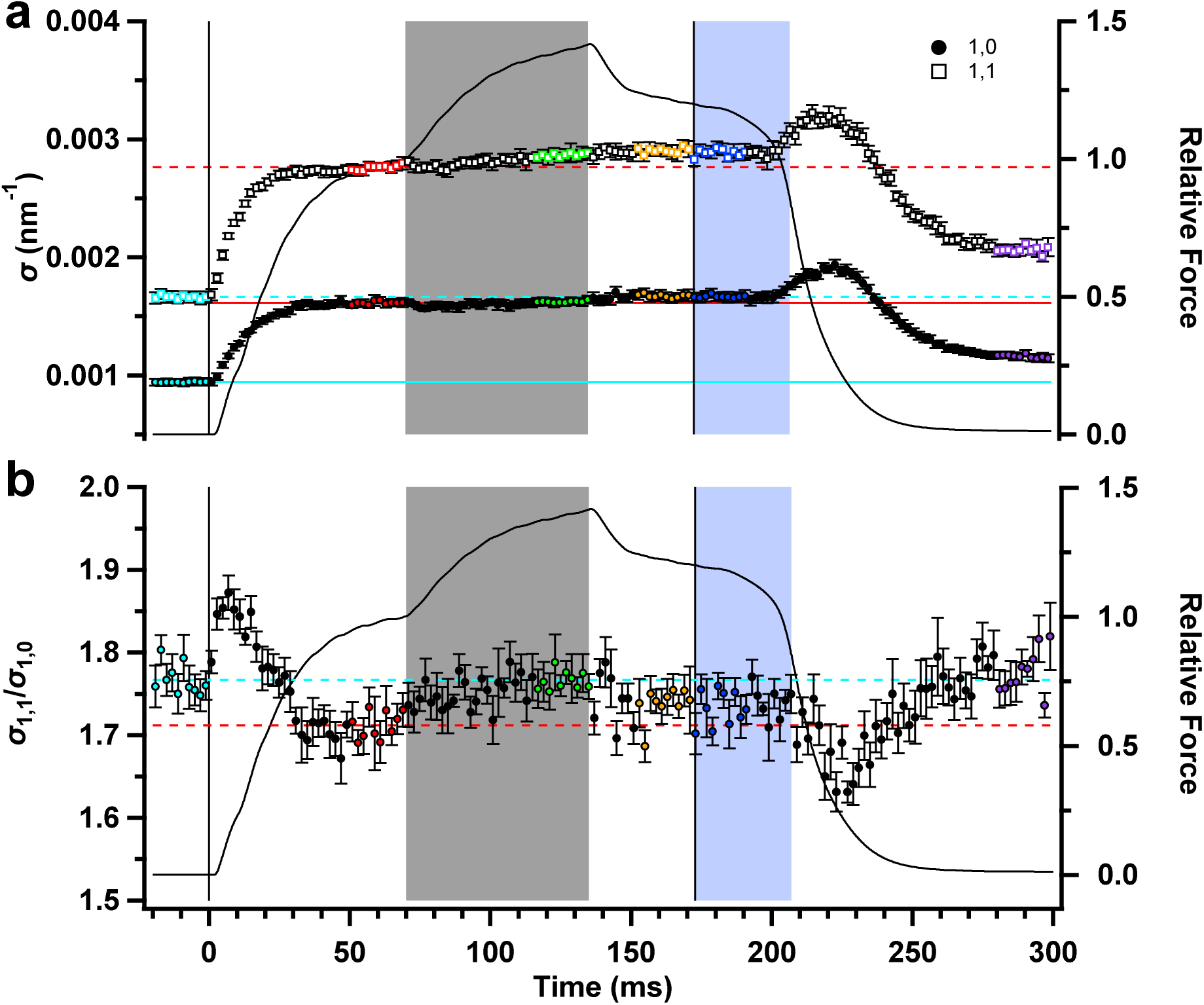
Radial widths of the equatorial reflections. **a** Radial width (σ) of the 1,0 (filled circles) and 1,1 (open squares) equatorial reflections. **b** Ratio of the radial widths of the 1,1 and 1,0 reflections (σ_1,1_/σ_1,0_). Colour coding of data points as in Fig. 1. Vertical solid lines denote the start and end of electrical stimulation; horizontal dashed lines denote the average resting value (cyan) and pre-stretch peak force value (red) of each parameter. Mean ± S.E.M. for n = 8 muscles.

**Fig. S3.**
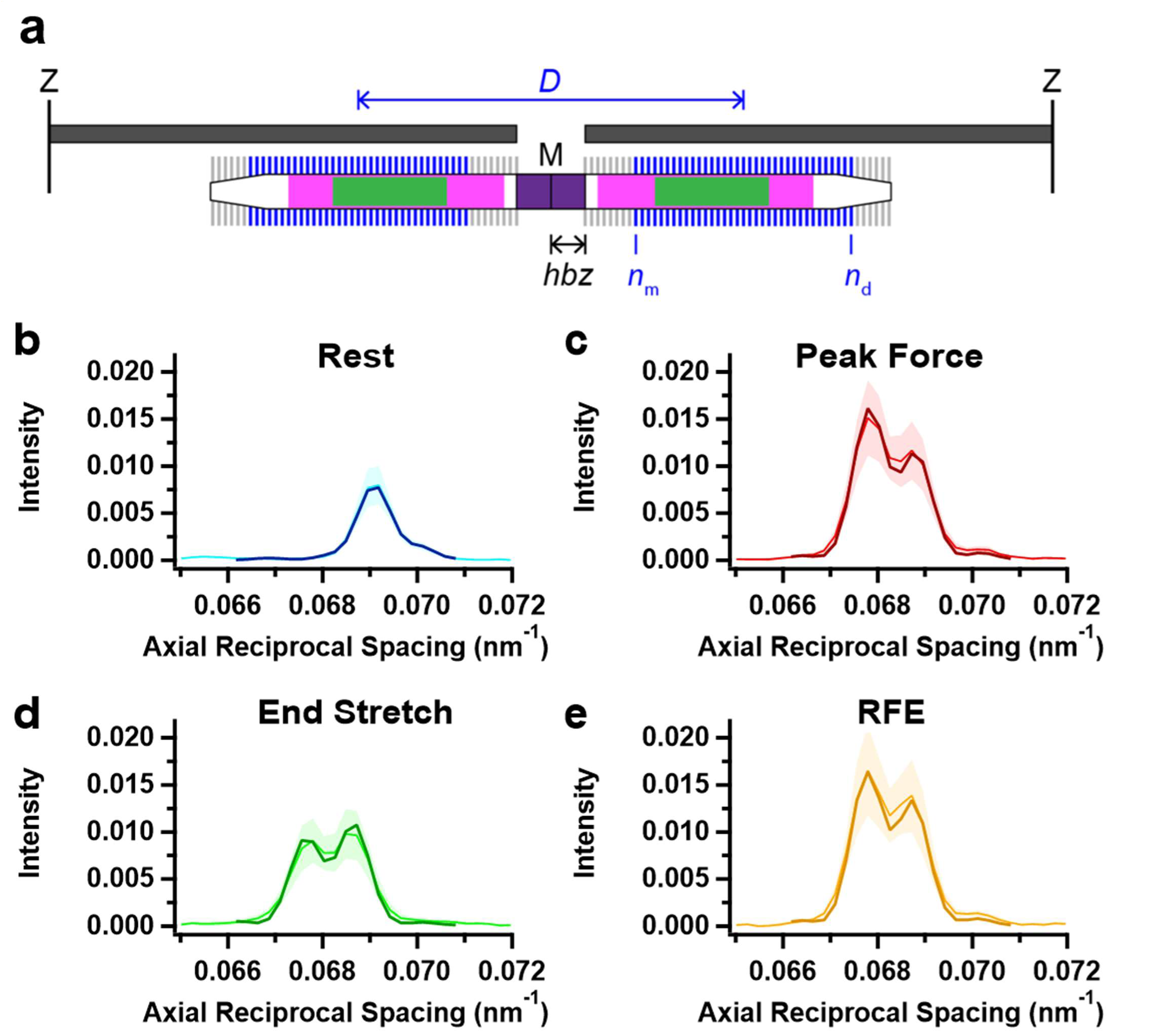
Fitting the axial profiles of the M3 reflection with a structural model of the thick filament. **a** Schematic diagram of the sarcomere showing model parameters. The sarcomere is delimited by Z-disks (black) and contains overlapping actin (dark grey) and myosin (white) filaments. Myosin filament midpoint, M; layers of myosin motors, vertical lines on myosin filament; bare zone at the centre of the sarcomere lacking myosin motors, purple; half-bare zone, *hbz*; zone of the thick filament containing titin C-type repeats, magenta; myosin-binding protein-C containing C-zone, green. Ordered layers of myosin motors (blue vertical bars between medial layer *n*_m_ and distal layer *n*_d_) in the two half-thick filaments have a centre-to-centre or interference distance *D* (blue). Disordered layers are shown in grey. **b** Experimental intensity distribution of the M3 at rest (cyan); **c** peak force before the ramp stretch (red); **d** the end of ramp stretch (green); **e** Residual force enhancement (RFE; orange). The best fit from the model in each case is superimposed as the darker colour in the reciprocal axial region 0.0662 – 0.0708 nm^−1^. Intensity distributions are the average of n=5 muscles at a camera length of 3.2m. Shaded areas around the experimental intensity distributions indicate ± S.D.

**Fig. S4.**
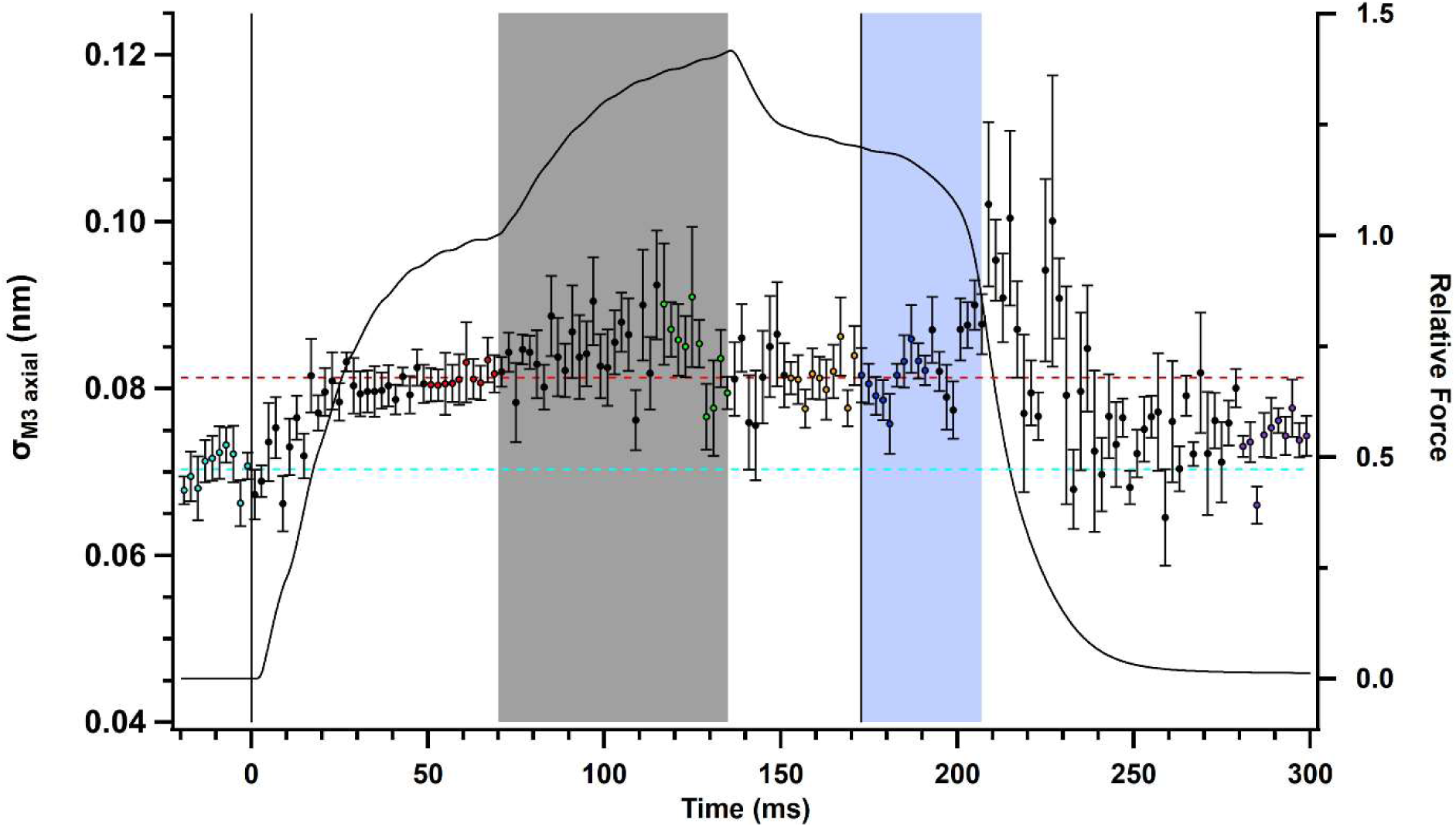
Axial width of the sub-peaks of the M3 reflection (σ_M3 axial_). Relative force (*T*/*T*_0_) is shown as the continuous black line. Colour coding of data points as in Fig. 1. Vertical solid lines denote the start and end of electrical stimulation; horizontal dashed lines denote the average resting value (cyan) and pre-stretch peak force value (red) of each parameter. Mean ± S.E.M. for n = 5 muscles for all data.

**Table S1.**
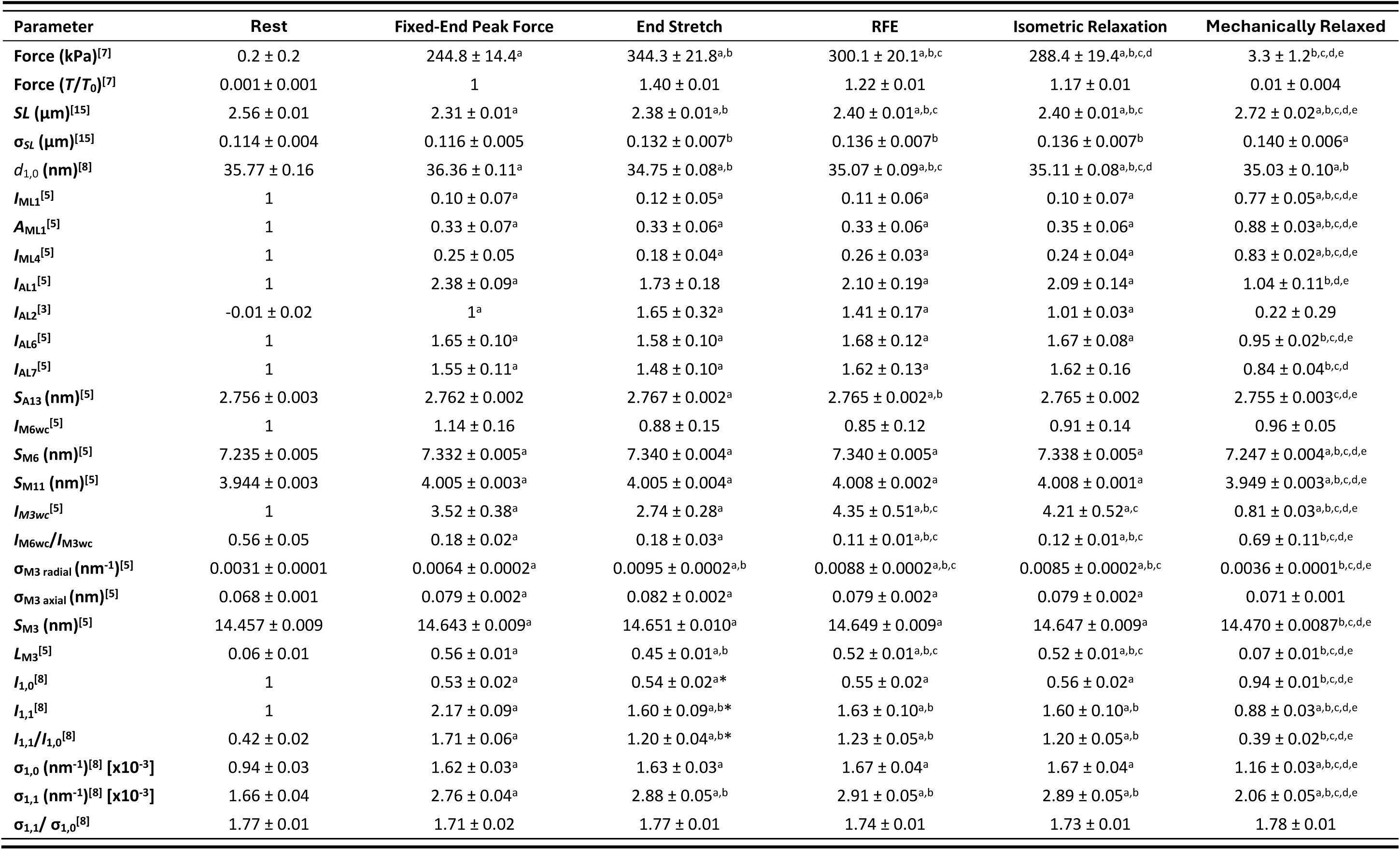
Change in force, sarcomere length and X-ray parameters in the periods defined in Fig 1a. Ten time frames were averaged from each muscle for each period. Data are presented as mean ± S.E.M. Superscripted letters denote significant differences between periods using a repeated-measures ANOVA with Tukey’s post hoc analyses. A significant main effect for the ANOVAs of each parameter was observed (P<0.001) except for *I*_M6_ where the main effect was P=0.185. Superscript letters denote significant differences between key phases. ^a^P<0.05 when compared to rest. ^b^P<0.05 when compared to peak force. ^c^P<0.05 when compared to end stretch. ^d^P<0.05 when compared to force enhanced state. ^e^P<0.05 when compared to isometric relaxation. ^[x]^number of technical repeats for each parameter.

**Table S2.**
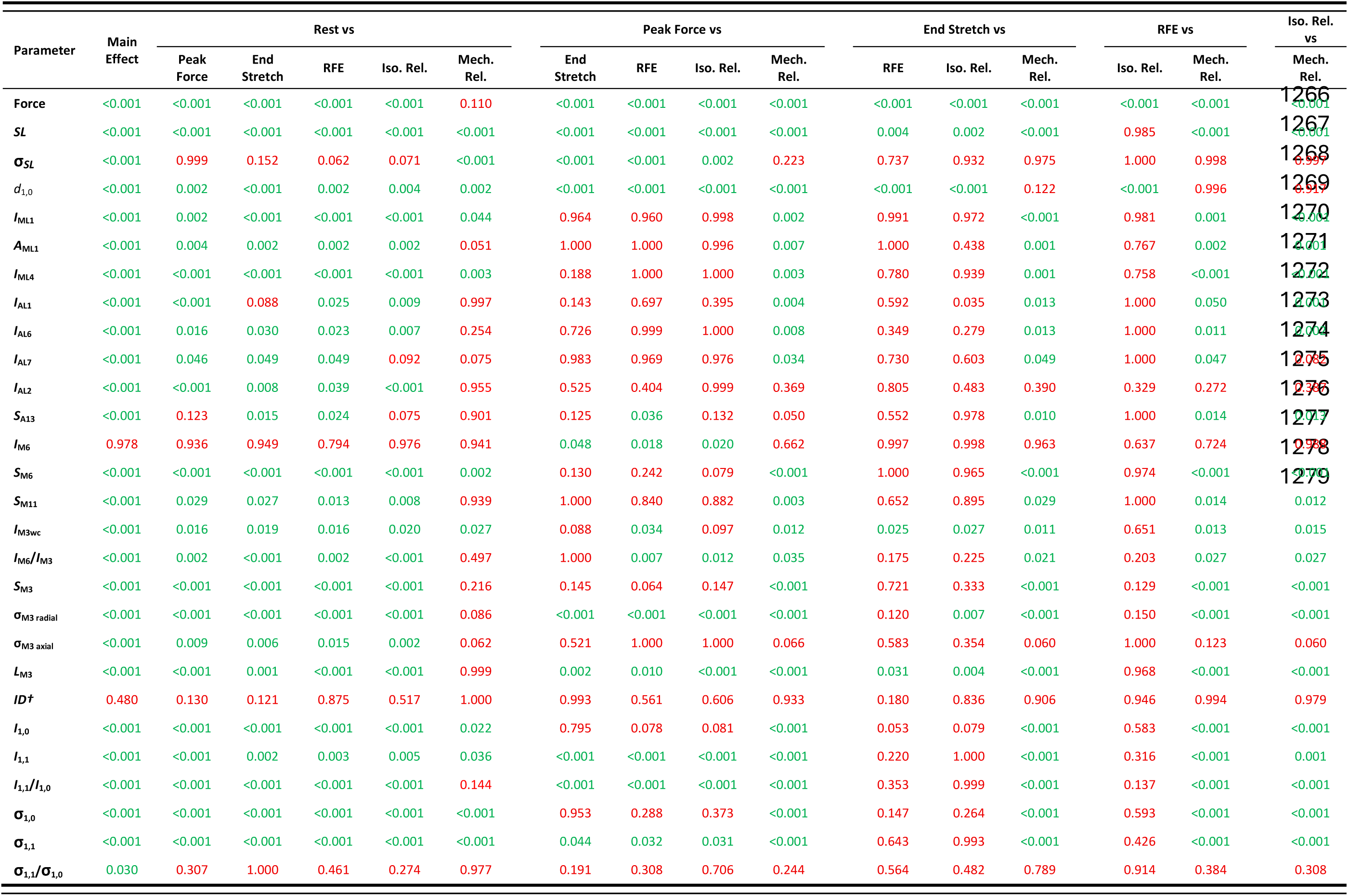
ANOVA output p-values.

## Notes

### Competing Interest Statement

The authors have declared no competing interest.

