## Supplementary figures and images for "The molecular structural basis of the braking action of muscle"

### Supplementary figures S1-S4

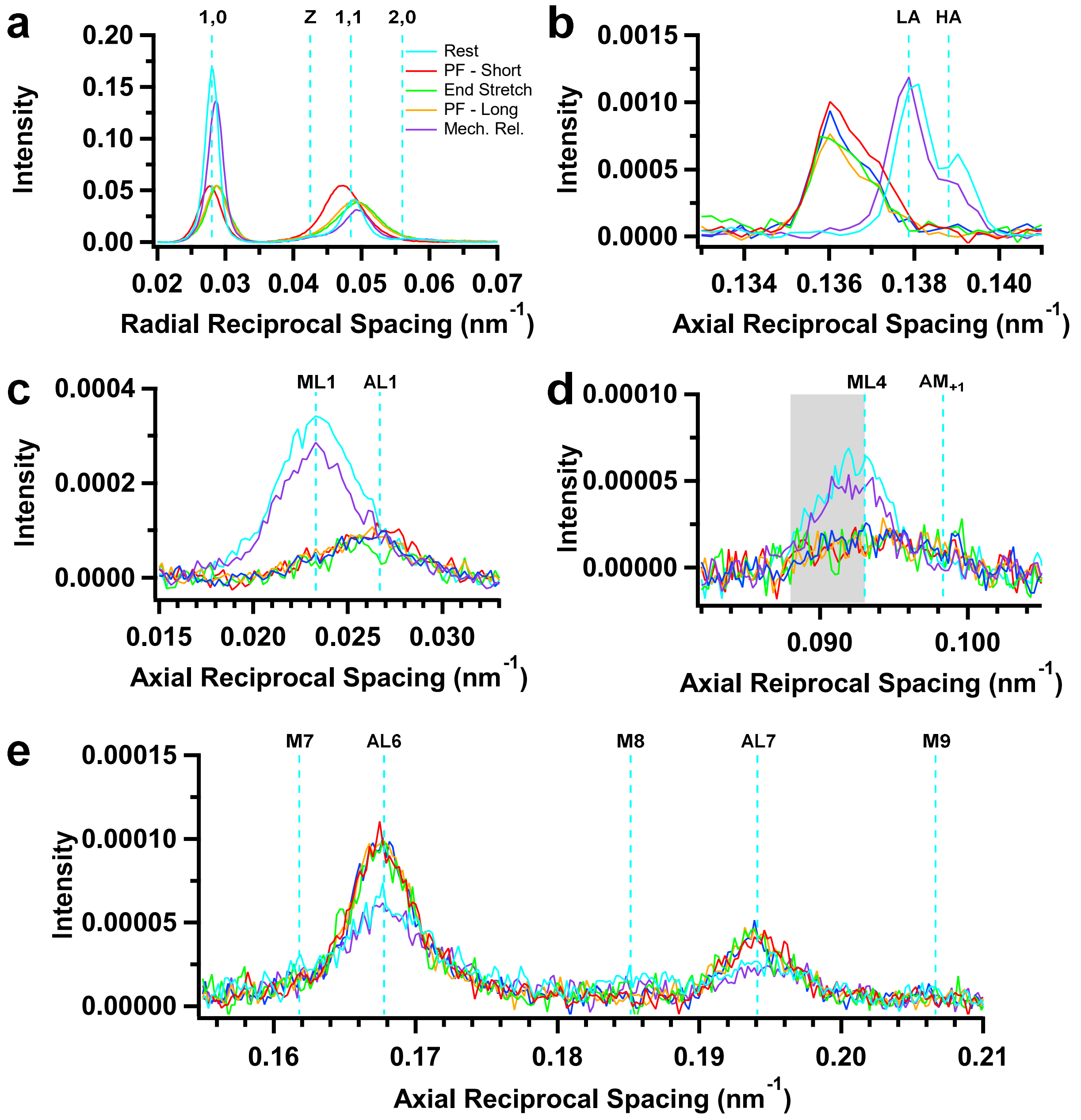

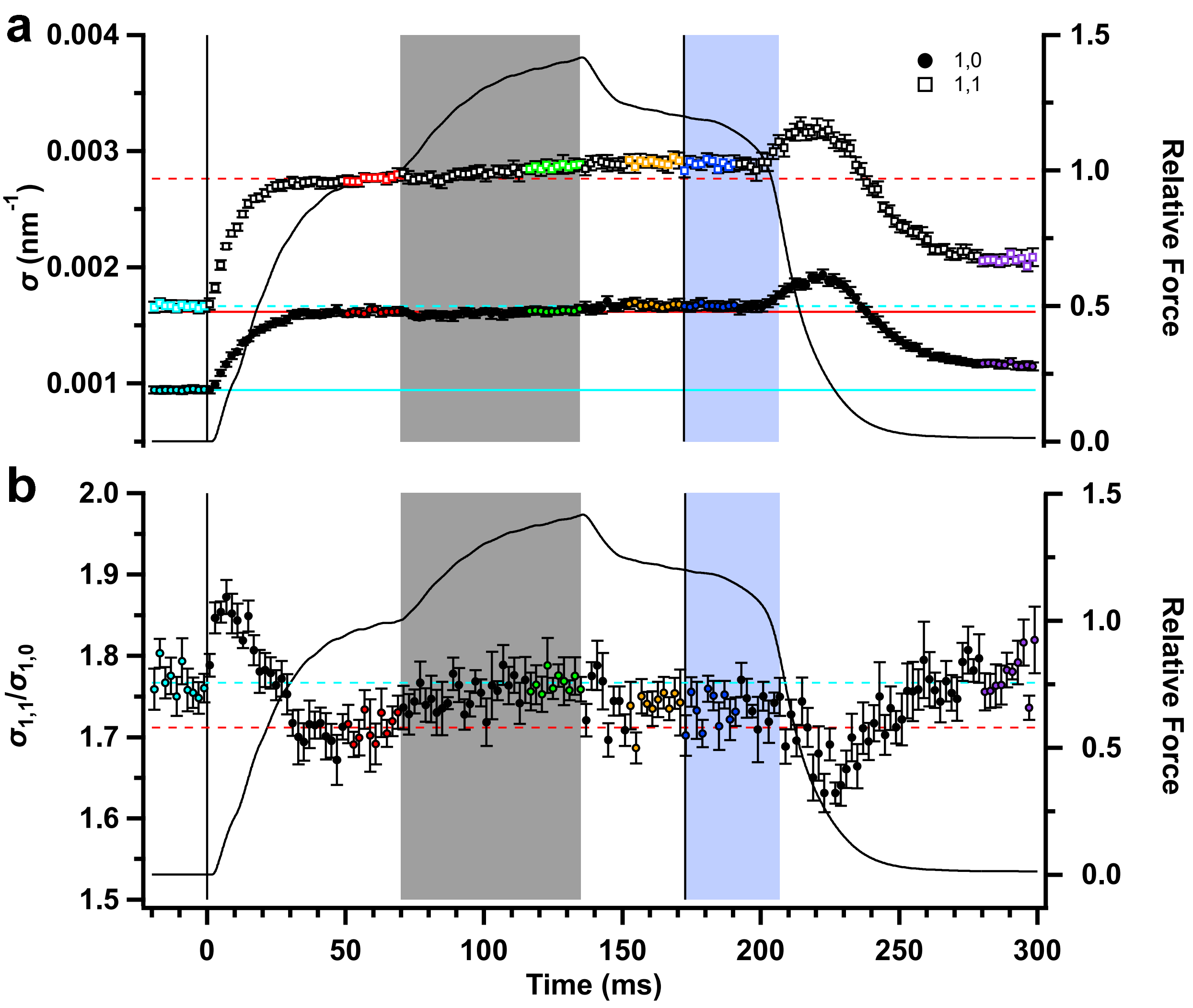

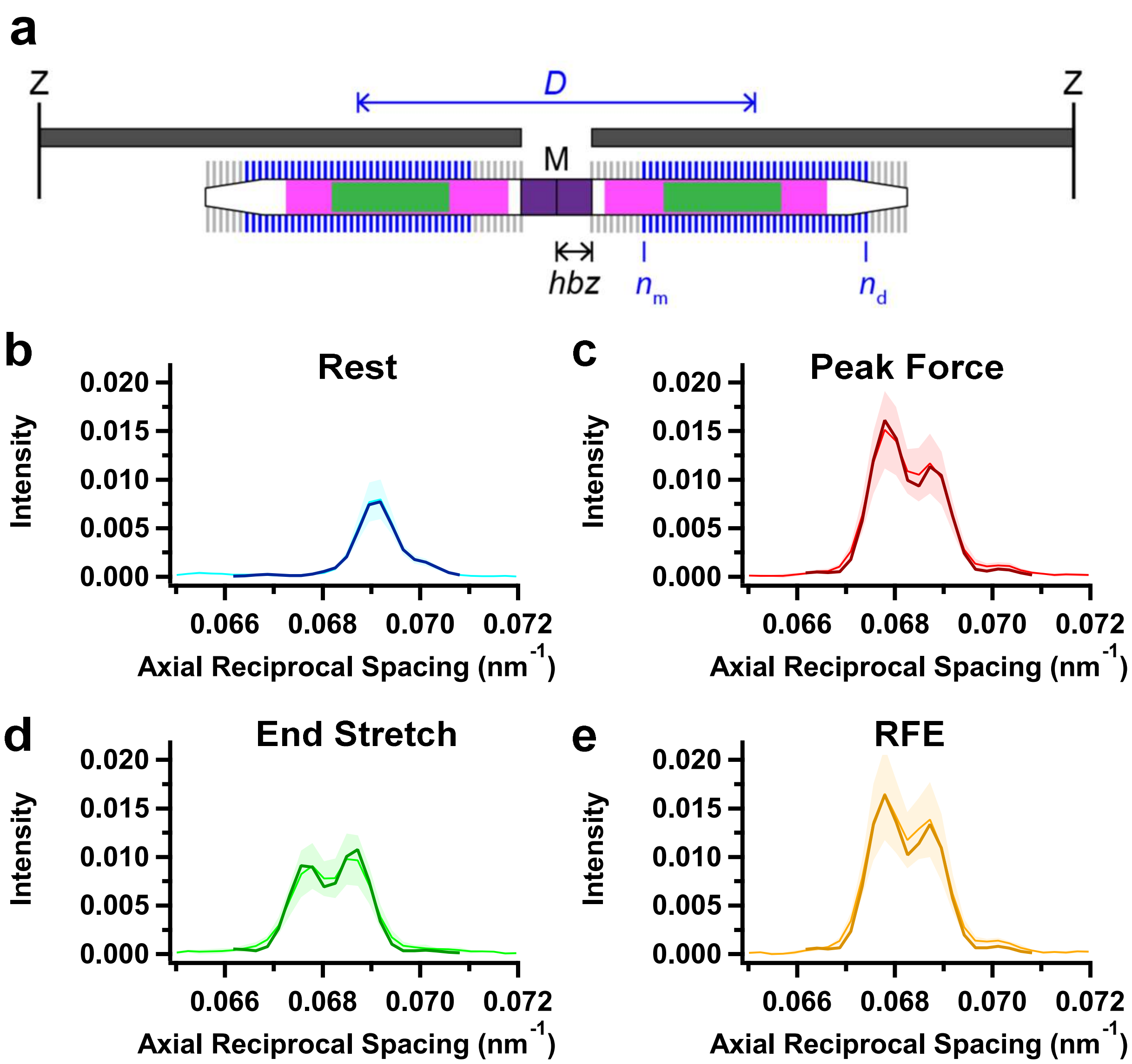

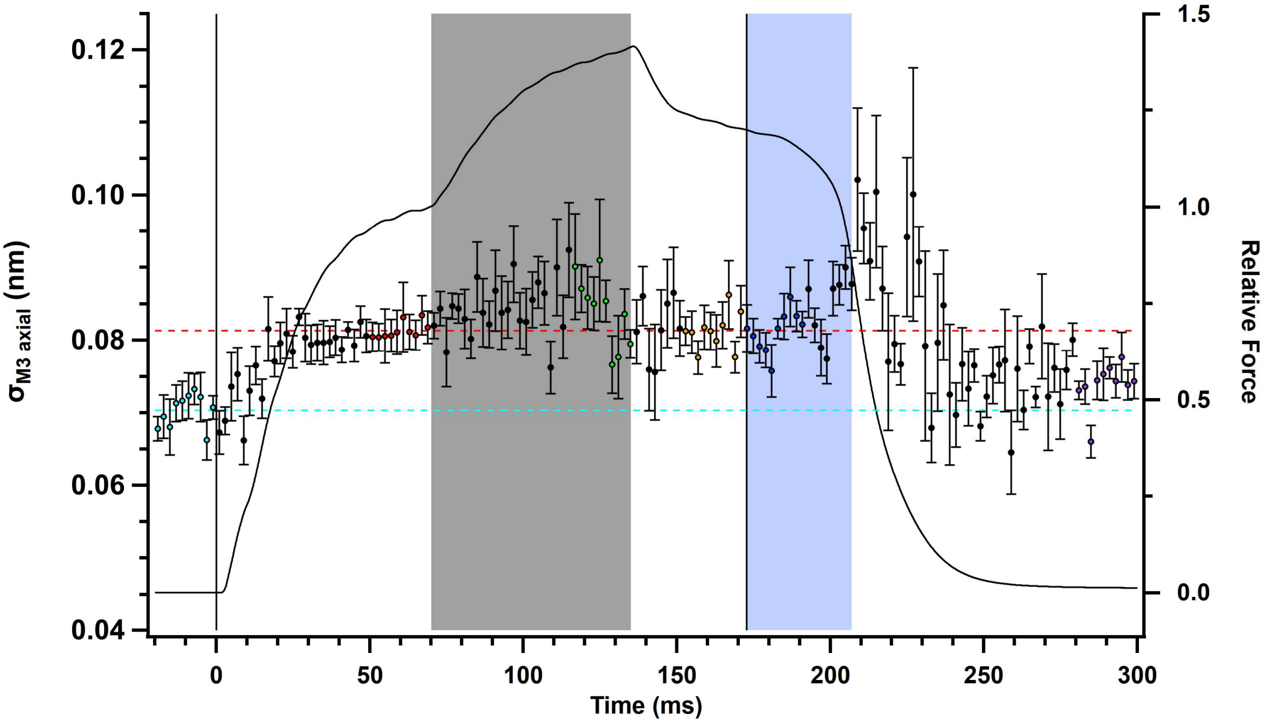
