## Supplementary tables S1 and S2 for "The molecular structural basis of the braking action of muscle"

| Parameter | Rest | Fixed-End Peak Force | End Stretch | RFE | Isometric Relaxation | Mechanically Relaxed |
|---|---|---|---|---|---|---|
| **Force (kPa)**[7] | 0.2 ± 0.2 | 244.8 ± 14.4[a] | 344.3 ± 21.8[a,b] | 300.1 ± 20.1[a,b,c] | 288.4 ± 19.4[a,b,c,d] | 3.3 ± 1.2[b,c,d,e] |
| **Force ($T/T_0$)**[7] | 0.001 ± 0.001 | 1 | 1.40 ± 0.01 | 1.22 ± 0.01 | 1.17 ± 0.01 | 0.01 ± 0.004 |
| **$SL$ (μm)**[15] | 2.56 ± 0.01 | 2.31 ± 0.01[a] | 2.38 ± 0.01[a,b] | 2.40 ± 0.01[a,b,c] | 2.40 ± 0.01[a,b,c] | 2.72 ± 0.02[a,b,c,d,e] |
| **$\sigma_{SL}$ (μm)**[15] | 0.114 ± 0.004 | 0.116 ± 0.005 | 0.132 ± 0.007[b] | 0.136 ± 0.007[b] | 0.136 ± 0.007[b] | 0.140 ± 0.006[a] |
| **$d_{1,0}$ (nm)**[8] | 35.77 ± 0.16 | 36.36 ± 0.11[a] | 34.75 ± 0.08[a,b] | 35.07 ± 0.09[a,b,c] | 35.11 ± 0.08[a,b,c,d] | 35.03 ± 0.10[a,b] |
| **$I_{ML1}$**[5] | 1 | 0.10 ± 0.07[a] | 0.12 ± 0.05[a] | 0.11 ± 0.06[a] | 0.10 ± 0.07[a] | 0.77 ± 0.05[a,b,c,d,e] |
| **$A_{ML1}$**[5] | 1 | 0.33 ± 0.07[a] | 0.33 ± 0.06[a] | 0.33 ± 0.06[a] | 0.35 ± 0.06[a] | 0.88 ± 0.03[a,b,c,d,e] |
| **$I_{ML4}$**[5] | 1 | 0.25 ± 0.05 | 0.18 ± 0.04[a] | 0.26 ± 0.03[a] | 0.24 ± 0.04[a] | 0.83 ± 0.02[a,b,c,d,e] |
| **$I_{AL1}$**[5] | 1 | 2.38 ± 0.09[a] | 1.73 ± 0.18 | 2.10 ± 0.19[a] | 2.09 ± 0.14[a] | 1.04 ± 0.11[b,d,e] |
| **$I_{AL2}$**[3] | -0.01 ± 0.02 | 1[a] | 1.65 ± 0.32[a] | 1.41 ± 0.17[a] | 1.01 ± 0.03[a] | 0.22 ± 0.29 |
| **$I_{AL6}$**[5] | 1 | 1.65 ± 0.10[a] | 1.58 ± 0.10[a] | 1.68 ± 0.12[a] | 1.67 ± 0.08[a] | 0.95 ± 0.02[b,c,d,e] |
| **$I_{AL7}$**[5] | 1 | 1.55 ± 0.11[a] | 1.48 ± 0.10[a] | 1.62 ± 0.13[a] | 1.62 ± 0.16 | 0.84 ± 0.04[b,c,d] |
| **$S_{A13}$ (nm)**[5] | 2.756 ± 0.003 | 2.762 ± 0.002 | 2.767 ± 0.002[a] | 2.765 ± 0.002[a,b] | 2.765 ± 0.002 | 2.755 ± 0.003[c,d,e] |
| **$I_{M6wc}$**[5] | 1 | 1.14 ± 0.16 | 0.88 ± 0.15 | 0.85 ± 0.12 | 0.91 ± 0.14 | 0.96 ± 0.05 |
| **$S_{M6}$ (nm)**[5] | 7.235 ± 0.005 | 7.332 ± 0.005[a] | 7.340 ± 0.004[a] | 7.340 ± 0.005[a] | 7.338 ± 0.005[a] | 7.247 ± 0.004[a,b,c,d,e] |
| **$S_{M11}$ (nm)**[5] | 3.944 ± 0.003 | 4.005 ± 0.003[a] | 4.005 ± 0.004[a] | 4.008 ± 0.002[a] | 4.008 ± 0.001[a] | 3.949 ± 0.003[a,b,c,d,e] |
| **$I_{M3wc}$**[5] | 1 | 3.52 ± 0.38[a] | 2.74 ± 0.28[a] | 4.35 ± 0.51[a,b,c] | 4.21 ± 0.52[a,c] | 0.81 ± 0.03[a,b,c,d,e] |
| **$I_{M6wc}/I_{M3wc}$** | 0.56 ± 0.05 | 0.18 ± 0.02[a] | 0.18 ± 0.03[a] | 0.11 ± 0.01[a,b,c] | 0.12 ± 0.01[a,b,c] | 0.69 ± 0.11[b,c,d,e] |
| **$\sigma_{M3\ radial}$ ($nm^{-1}$)**[5] | 0.0031 ± 0.0001 | 0.0064 ± 0.0002[a] | 0.0095 ± 0.0002[a,b] | 0.0088 ± 0.0002[a,b,c] | 0.0085 ± 0.0002[a,b,c] | 0.0036 ± 0.0001[b,c,d,e] |
| **$\sigma_{M3\ axial}$ (nm)**[5] | 0.068 ± 0.001 | 0.079 ± 0.002[a] | 0.082 ± 0.002[a] | 0.079 ± 0.002[a] | 0.079 ± 0.002[a] | 0.071 ± 0.001 |
| **$S_{M3}$ (nm)**[5] | 14.457 ± 0.009 | 14.643 ± 0.009[a] | 14.651 ± 0.010[a] | 14.649 ± 0.009[a] | 14.647 ± 0.009[a] | 14.470 ± 0.0087[b,c,d,e] |
| **$L_{M3}$**[5] | 0.06 ± 0.01 | 0.56 ± 0.01[a] | 0.45 ± 0.01[a,b] | 0.52 ± 0.01[a,b,c] | 0.52 ± 0.01[a,b,c] | 0.07 ± 0.01[b,c,d,e] |
| **$I_{1,0}$**[8] | 1 | 0.53 ± 0.02[a] | 0.54 ± 0.02[a]* | 0.55 ± 0.02[a] | 0.56 ± 0.02[a] | 0.94 ± 0.01[b,c,d,e] |
| **$I_{1,1}$**[8] | 1 | 2.17 ± 0.09[a] | 1.60 ± 0.09[a,b]* | 1.63 ± 0.10[a,b] | 1.60 ± 0.10[a,b] | 0.88 ± 0.03[a,b,c,d,e] |
| **$I_{1,1}/I_{1,0}$**[8] | 0.42 ± 0.02 | 1.71 ± 0.06[a] | 1.20 ± 0.04[a,b]* | 1.23 ± 0.05[a,b] | 1.20 ± 0.05[a,b] | 0.39 ± 0.02[b,c,d,e] |
| **$\sigma_{1,0}$ ($nm^{-1}$)**[8] **[$x10^{-3}$]** | 0.94 ± 0.03 | 1.62 ± 0.03[a] | 1.63 ± 0.03[a] | 1.67 ± 0.04[a] | 1.67 ± 0.04[a] | 1.16 ± 0.03[a,b,c,d,e] |
| **$\sigma_{1,1}$ ($nm^{-1}$)**[8] **[$x10^{-3}$]** | 1.66 ± 0.04 | 2.76 ± 0.04[a] | 2.88 ± 0.05[a,b] | 2.91 ± 0.05[a,b] | 2.89 ± 0.05[a,b] | 2.06 ± 0.05[a,b,c,d,e] |
| **$\sigma_{1,1}/\sigma_{1,0}$**[8] | 1.77 ± 0.01 | 1.71 ± 0.02 | 1.77 ± 0.01 | 1.74 ± 0.01 | 1.73 ± 0.01 | 1.78 ± 0.01 |

**Table S1 | Change in force, sarcomere length and X-ray parameters in the periods defined in Fig 1a.** Ten time frames were averaged from each muscle for each period. Data are presented as mean ± S.E.M. Superscripted letters denote significant differences between periods using a repeated-measures ANOVA with Tukey's post hoc analyses. A significant main effect for the ANOVAs of each parameter was observed ($P<0.001$) except for $I_{M6}$ where the main effect was $P=0.185$. Superscript letters denote significant differences between key phases. [a]P<0.05 when compared to rest. [b]P<0.05 when compared to peak force. [c]P<0.05 when compared to end stretch. [d]P<0.05 when compared to force enhanced state. [e]P<0.05 when compared to isometric relaxation. [x]number of technical repeats for each parameter.

| Parameter | Main Effect | Rest vs | | | | | Peak Force vs | | | | End Stretch vs | | | RFE vs | | Iso. Rel. vs |
|---|---|---|---|---|---|---|---|---|---|---|---|---|---|---|---|---|
| | | Peak Force | End Stretch | RFE | Iso. Rel. | Mech. Rel. | End Stretch | RFE | Iso. Rel. | Mech. Rel. | RFE | Iso. Rel. | Mech. Rel. | Iso. Rel. | Mech. Rel. | Mech. Rel. |
| **Force** | <0.001 | <0.001 | <0.001 | <0.001 | <0.001 | 0.110 | <0.001 | <0.001 | <0.001 | <0.001 | <0.001 | <0.001 | <0.001 | <0.001 | <0.001 | <0.001 |
| $SL$ | <0.001 | <0.001 | <0.001 | <0.001 | <0.001 | <0.001 | <0.001 | <0.001 | <0.001 | <0.001 | 0.004 | 0.002 | <0.001 | 0.985 | <0.001 | <0.001 |
| $\sigma_{SL}$ | <0.001 | 0.999 | 0.152 | 0.062 | 0.071 | <0.001 | <0.001 | <0.001 | 0.002 | 0.223 | 0.737 | 0.932 | 0.975 | 1.000 | 0.998 | 0.997 |
| $d_{1,0}$ | <0.001 | 0.002 | <0.001 | 0.002 | 0.004 | 0.002 | <0.001 | <0.001 | <0.001 | <0.001 | <0.001 | <0.001 | 0.122 | <0.001 | 0.996 | 0.917 |
| $I_{ML1}$ | <0.001 | 0.002 | <0.001 | <0.001 | <0.001 | 0.044 | 0.964 | 0.960 | 0.998 | 0.002 | 0.991 | 0.972 | <0.001 | 0.981 | 0.001 | <0.001 |
| $A_{ML1}$ | <0.001 | 0.004 | 0.002 | 0.002 | 0.002 | 0.051 | 1.000 | 1.000 | 0.996 | 0.007 | 1.000 | 0.438 | 0.001 | 0.767 | 0.002 | 0.001 |
| $I_{ML4}$ | <0.001 | <0.001 | <0.001 | <0.001 | <0.001 | 0.003 | 0.188 | 1.000 | 1.000 | 0.003 | 0.780 | 0.939 | 0.001 | 0.758 | <0.001 | <0.001 |
| $I_{AL1}$ | <0.001 | <0.001 | 0.088 | 0.025 | 0.009 | 0.997 | 0.143 | 0.697 | 0.395 | 0.004 | 0.592 | 0.035 | 0.013 | 1.000 | 0.050 | 0.001 |
| $I_{AL6}$ | <0.001 | 0.016 | 0.030 | 0.023 | 0.007 | 0.254 | 0.726 | 0.999 | 1.000 | 0.008 | 0.349 | 0.279 | 0.013 | 1.000 | 0.011 | 0.002 |
| $I_{AL7}$ | <0.001 | 0.046 | 0.049 | 0.049 | 0.092 | 0.075 | 0.983 | 0.969 | 0.976 | 0.034 | 0.730 | 0.603 | 0.049 | 1.000 | 0.047 | 0.082 |
| $I_{AL2}$ | <0.001 | <0.001 | 0.008 | 0.039 | <0.001 | 0.955 | 0.525 | 0.404 | 0.999 | 0.369 | 0.805 | 0.483 | 0.390 | 0.329 | 0.272 | 0.387 |
| $S_{A13}$ | <0.001 | 0.123 | 0.015 | 0.024 | 0.075 | 0.901 | 0.125 | 0.036 | 0.132 | 0.050 | 0.552 | 0.978 | 0.010 | 1.000 | 0.014 | 0.013 |
| $I_{M6}$ | 0.978 | 0.936 | 0.949 | 0.794 | 0.976 | 0.941 | 0.048 | 0.018 | 0.020 | 0.662 | 0.997 | 0.998 | 0.963 | 0.637 | 0.724 | 0.988 |
| $S_{M6}$ | <0.001 | <0.001 | <0.001 | <0.001 | <0.001 | 0.002 | 0.130 | 0.242 | 0.079 | <0.001 | 1.000 | 0.965 | <0.001 | 0.974 | <0.001 | <0.001 |
| $S_{M11}$ | <0.001 | 0.029 | 0.027 | 0.013 | 0.008 | 0.939 | 1.000 | 0.840 | 0.882 | 0.003 | 0.652 | 0.895 | 0.029 | 1.000 | 0.014 | 0.012 |
| $I_{M3wc}$ | <0.001 | 0.016 | 0.019 | 0.016 | 0.020 | 0.027 | 0.088 | 0.034 | 0.097 | 0.012 | 0.025 | 0.027 | 0.011 | 0.651 | 0.013 | 0.015 |
| $I_{M6}/I_{M3}$ | <0.001 | 0.002 | <0.001 | 0.002 | <0.001 | 0.497 | 1.000 | 0.007 | 0.012 | 0.035 | 0.175 | 0.225 | 0.021 | 0.203 | 0.027 | 0.027 |
| $S_{M3}$ | <0.001 | <0.001 | <0.001 | <0.001 | <0.001 | 0.216 | 0.145 | 0.064 | 0.147 | <0.001 | 0.721 | 0.333 | <0.001 | 0.129 | <0.001 | <0.001 |
| $\sigma_{M3\ radial}$ | <0.001 | <0.001 | <0.001 | <0.001 | <0.001 | 0.086 | <0.001 | <0.001 | <0.001 | <0.001 | 0.120 | 0.007 | <0.001 | 0.150 | <0.001 | <0.001 |
| $\sigma_{M3\ axial}$ | <0.001 | 0.009 | 0.006 | 0.015 | 0.002 | 0.062 | 0.521 | 1.000 | 1.000 | 0.066 | 0.583 | 0.354 | 0.060 | 1.000 | 0.123 | 0.060 |
| $L_{M3}$ | <0.001 | <0.001 | 0.001 | <0.001 | <0.001 | 0.999 | 0.002 | 0.010 | <0.001 | <0.001 | 0.031 | 0.004 | <0.001 | 0.968 | <0.001 | <0.001 |
| ***ID†*** | 0.480 | 0.130 | 0.121 | 0.875 | 0.517 | 1.000 | 0.993 | 0.561 | 0.606 | 0.933 | 0.180 | 0.836 | 0.906 | 0.946 | 0.994 | 0.979 |
| $I_{1,0}$ | <0.001 | <0.001 | <0.001 | <0.001 | <0.001 | 0.022 | 0.795 | 0.078 | 0.081 | <0.001 | 0.053 | 0.079 | <0.001 | 0.583 | <0.001 | <0.001 |
| $I_{1,1}$ | <0.001 | <0.001 | 0.002 | 0.003 | 0.005 | 0.036 | <0.001 | <0.001 | <0.001 | <0.001 | 0.220 | 1.000 | <0.001 | 0.316 | <0.001 | 0.001 |
| $I_{1,1}/I_{1,0}$ | <0.001 | <0.001 | <0.001 | <0.001 | <0.001 | 0.144 | <0.001 | <0.001 | <0.001 | <0.001 | 0.353 | 0.999 | <0.001 | 0.137 | <0.001 | <0.001 |
| $\sigma_{1,0}$ | <0.001 | <0.001 | <0.001 | <0.001 | <0.001 | <0.001 | 0.953 | 0.288 | 0.373 | <0.001 | 0.147 | 0.264 | <0.001 | 0.593 | <0.001 | <0.001 |
| $\sigma_{1,1}$ | <0.001 | <0.001 | <0.001 | <0.001 | <0.001 | <0.001 | 0.044 | 0.032 | 0.031 | <0.001 | 0.643 | 0.993 | <0.001 | 0.426 | <0.001 | <0.001 |
| $\sigma_{1,1}/\sigma_{1,0}$ | 0.030 | 0.307 | 1.000 | 0.461 | 0.274 | 0.977 | 0.191 | 0.308 | 0.706 | 0.244 | 0.564 | 0.482 | 0.789 | 0.914 | 0.384 | 0.308 |

**Table S2 | ANOVA output p-values.**
